# Genome-Wide Association study in a US soft winter wheat population reveals novel and known sources of resistance to the Septoria tritici blotch pathogen *Zymoseptoria tritici*

**DOI:** 10.1101/2025.09.30.679614

**Authors:** Lamia Aouini, Rupesh Gaire, Steven Scofield, Gina Brown-Guedira, Mohsen Mohammadi, Stephen B. Goodwin

## Abstract

*Key message* GWAS of 337 soft winter wheat genotypes from a breeding program in Indiana, USA identified marker-trait associations that likely correspond with existing and novel resistances to Septoria tritici blotch disease.

**Abstract** Septoria tritici blotch (STB), caused by the ascomycete fungus *Zymoseptoria tritici*, is a major disease of wheat worldwide. To find additional sources of resistance, Genome-Wide Association (GWA) was used to analyze 337 soft winter wheat genotypes from Indiana, USA, by inoculating seedlings with two isolates of *Z. tritici* in a complete randomized design. Necrosis and pycnidia development were assessed at 14, 18 and 22 days post inoculation, enabling the calculation of area under the disease progress curve (AUDPC) for each parameter. Adjusted necrosis and pycnidia AUDPC scores were compared with 14,341 high-quality SNPs in a GWA analysis using the FarmCPU and GAPIT CMLM models to identify markers associated with the resistance. Significant (p < 0.05) isolate × genotype interactions were identified, confirming that the phenotypic variation was caused by isolate-specific resistance genes. Overall, 12 marker-trait associations (MTAs) were identified with *Z. tritici* necrosis and pycnidia resistance. All mapped MTAs were isolate and necrosis/pycnidia specific. Distinct MTAs were mapped for necrosis and pycnidia on chromosome 6A, and for pycnidia on chromosomes 1A, 1D, 3B, 4A, 5B, 6A and 7A. MTAs on chromosomes 5B, 4A, 6A, and 1D likely corresponded with the known genes *Stb1*, *Stb7*, *Stb15* and *Stb19*, respectively. MTAs on chromosomes 1A, 3B, 4A, 6A and 7A were not associated with previously known *Stb* genes and may be novel. Candidate genes near the marker locations have been identified for further investigations. The GWAS showed that Indiana soft winter wheat germplasm segregates for novel and known *Stb* genes and constitutes a valuable breeding resource for *Z. tritici* resistance.

## Introduction

Wheat is a universal crop that has facilitated the dramatic shift from hunter-gatherer to agrarian societies during the evolution of human civilization (Eckardt 2010). Since its domestication 10,000 years ago, wheat has contributed greatly to human nutrition and is currently a leading source of calories and plant-derived protein, contributing up to 35% of global dietary intake (Ali et al. 2020; Shewry 2009). Tremendous breeding efforts have been made to meet the ever-increasing human population and concomitant rise in demand for food (Khush 2001). These efforts have led to an unprecedented increase in wheat production and a steady balance of supply versus demand (Figueroa et al. 2018). Nonetheless, feeding the growing global population, predicted to exceed nine billion people by 2050, will require an estimated annual yield increase of one billion tonnes. Moreover, fast-tracking climate shifts that promote the emergence of increasingly unpredictable biotic and abiotic pressures are expanding threats to wheat production that must be considered when breeding for superior wheat cultivars (Fones et al. 2020).

Fungal pathogens are among the most perilous biotic threats to wheat production (Figueroa et al. 2018; Seybold et al. 2020). *Zymoseptoria tritici* (formerly known as *Mycosphaerella graminicola* and *Septoria tritici*), the causal agent of Septoria tritici blotch (STB), is a devastating pathogen (Fones et al. 2020; Seybold et al. 2020) that accounts for average yield losses of 5 to 10% per year in the main wheat-producing countries, including the U.S. (Cowger et al. 2020; Fones and Gurr 2015). Despite its leading position in soft winter wheat with an annual production of 15.5 million tons in 2019 (NASS 2020), U.S. wheat crops are constrained by frequent attacks by *Z. tritici*. STB has been among the most destructive wheat diseases in the Midwestern state of Indiana since at least 1955 (Shaner and Buechley 1995). Thus far, STB has been controlled mostly with fungicides, rendering it the major target for the entire agrochemical industry (Cools and Fraaije 2008; Lendenmann et al. 2015; Lucas et al. 2015; Torriani et al. 2015). However, increasing issues with the numbers of fungicide applications, and resistance phenomena in the fungus, underscore the necessity for more sustainable approaches to disease management (Cheval et al. 2017; Heick et al. 2017; Lendenmann et al. 2015; Mundt et al. 2002; Omrane et al. 2015; Price et al. 2015; Torriani et al. 2015).

Host resistance is a sustainable and potentially effective alternative to limit the devastating effects of this perilous pathogen. In Indiana, efforts began during the 1960s to breed for resistant cultivars of soft winter wheat (Shaner and Finney 1982). These efforts led to the identification of Bulgaria 88 (PI94407), which carries the major resistance gene *Stb1* (Adhikari et al. 2004a; Shaner and Finney 1982). This gene was transferred from Bulgaria 88 into soft red winter wheat cultivars Oasis and Sullivan, which had superior agronomic qualities and were first released during 1975 and 1979, respectively (Adhikari et al. 2004a). The major gene *Stb1* has proven its efficacy in diverse STB-prone environments in the U.S and across the world over many years and therefore appears to be durable (Adhikari et al. 2004a; Brown et al. 2015a). This success, coupled with attention brought by damaging STB epidemics such as one that occurred in North Africa between 1968 and 1969 (Saadaoui 1987), has spurred international awareness and breeding efforts. This work plus an increased focus on genetic analysis has led to the discovery of at least 22 major resistance genes and more than 170 quantitative trait loci (QTL) for enhanced *Z. tritici* resistance in wheat, mainly identified through analyses of bi-parental mapping populations (Brown et al. 2015a; Langlands-Perry et al. 2022; Yang et al. 2018; 2021).

Resistance to *Z. tritici* in wheat is a complex trait that has been investigated intensively (Brown et al. 2015a; Ghaffary 2011; Ghaffary et al. 2018; Goodwin 2007; Gurung et al. 2014). Despite the success of linkage mapping studies in analyzing multiple qualitative and quantitative trait loci, the genetic complexity of STB resistance has hampered an accurate understanding of the phenotypic variation, and thus an effective translation to STB breeding (Mirdita et al. 2015). There remains a need for improved mapping resolution power to detect variable loci, especially in cases where the underlying genetic architecture is highly polygenic and complex (Corwin and Kliebenstein 2017).

This need for identification of additional loci for STB resistance can be satisfied by genome-wide association studies (GWAS), which have been used intensively to decode complex phenotypic traits and to identify genomic regions associated with variation in broad-based and diverse populations of many species (Liu and Yan 2019). Originally, association studies were adopted from human medical genetics, and were first deployed in plants during 2001 to dissect sequence polymorphisms in the *Dwarf8* gene of maize (Thornsberry et al. 2001). Ever since, GWA studies have proliferated and have been adopted intensively to dissect the genetic architecture of multiple quantitative traits in many commercial crops (Alqudah et al. 2020; Jabbari et al. 2018; Pantaliao et al. 2016; Razifard et al. 2020), including wheat (Battenfield et al. 2016; Lopes et al. 2015; Neumann et al. 2011; Singh et al. 2018; Sukumaran et al. 2015; Tsai et al. 2020), where GWAS provided a valuable tool to find molecular markers related to a number of fungal diseases including rust (Juliana et al. 2018; Ledesma-Ramírez et al. 2019; Maccaferri et al. 2010; Prins et al. 2016; Yu et al. 2012), tan spot (Galagedara et al. 2020; Phuke et al. 2020), Fusarium head blight (Miedaner et al. 2011) and STB (Ballini et al. 2020; Kristensen et al. 2018; Mikaberidze et al 2025; Muqaddasi et al. 2019b).

A few GWA studies have been targeted at identifying the associations between genomic regions of wheat and resistance to *Z. tritici* in elite cultivars and landraces (Ando et al. 2018; Kidane et al. 2017; Kollers et al. 2013; Mekonnen et al. 2021; Miedaner et al. 2013; Odilbekov et al. 2019; Yang et al. 2022). For instance, four significant single-nucleotide polymorphisms (SNPs) were identified in a study by Miedaner et al. (2013) and were associated with STB resistance located on chromosomes 1B, 2B, 5B, and 6A. That analysis was conducted on 1,055 elite hybrids and their corresponding 87 parental lines. Other studies have used hexaploid wheat landraces (Odilbekov et al. 2019) or breeding lines (Mekonen et al. 2021; Vagndorf et al. 2017; Yang et al. 2022) in a GWA approach to detect genomic regions associated with STB resistance. The previous GWA analyses of bread wheat cultivars were performed on populations from Australia, Europe, the northwestern United States or diverse international sources; no comparable analyses have been performed on soft, red, winter wheat cultivars from the Midwestern U.S., which may be more likely to identify novel resistance alleles that would be effective against populations of *Z. tritici* in that region of North America.

The goal of this research was to assay a population of 337 soft winter wheat lines in a GWA analysis to test the hypothesis that wheat lines and cultivars selected in Indiana, USA would contain novel sources of seedling resistance against STB caused by *Z. tritici*. Specific objectives to achieve this goal were to (1) identify markers associated with *Z. tritici* resistance; (2) compare two GWA models for a better and powerful mapping resolution; (3) test whether analyses at a single time point would give the same results as calculations based on Area Under the Disease Progress Curve using multiple scoring times; (4) compare the obtained results with the chromosomal locations of known *Stb* resistance genes and QTL; and (5) identify candidate STB genes in the soft winter wheat population.

## Materials and methods

### Plant materials and *Zymoseptoria tritici* isolates

Screening for STB resistance was performed with three sets of hexaploid wheat lines. The first and second sets were comprised of 11 and 21 wheat lines, respectively, with known and predicted STB resistance (Table 1). The remaining set, the primary object of this analysis, was a population of 337 soft winter wheat lines from the Purdue University small grains breeding program derived from 123 crosses conducted between 1988 and 2010 (Gaire et al. 2020), designated hereafter as the “soft red winter wheat panel” population.

**Table 1.** Hexaploid wheat lines screened for levels of Septoria tritici blotch with diverse isolates of *Zymoseptoria tritici*.

| Cultivar or line | Growth habit<sup>a</sup> | Origin | Stb gene (chromosome arm) | Experiment<sup>b</sup> | Reference |
| --- | --- | --- | --- | --- | --- |
| Bulgaria 88<sup>c</sup> | W | Bulgaria | *Stb1* (5BL) | 2 | Adhikari et al. 2004b |
| Oasis | W | USA | *Stb1* (5BL) | 1,2 | Adhikari et al. 2004b |
| Veranopolis<sup>c</sup> | S | Brazil | *Stb2* (3BS) | 1,2 | Liu et al. 2013 |
| Israel 493<sup>c</sup> | S | Israel | *Stb3* (7AS) | 2 | Goodwin et al. 2015 |
| Tadinia<sup>c</sup> | S | USA | *Stb4* (7DS) | 1,2 | Adhikari et al. 2004a |
| Cs Synthetic (6x)7D<sup>c</sup> | S | China / USA | *Stb5* (7DS) | 2 | Arraiano et al. 2001 |
| Shafir | S | Israel | *Stb6* (3AS) | 2 | Chartrain et al. 2005b |
| Estanzuela Federal | S | Uruguay | *Stb7* (4AL) | 2 | McCartney et al. 2003 |
| M6/Synthetic (W7984) | W | USA | *Stb8* (7BL) | 2 | Adhikari et al. 2003 |
| Courtot | W | France | *Stb9* (2BL) | 2 | Chartrain et al. 2009 |
| Kavkaz-K4500<sup>c,d</sup> | F | CIMMYT | *Stb10* (1D) + *Stb12* (4AL) | 2 | Chartrain et al. 2005a |
| TE9111<sup>c,d</sup> | S | Portugal | *Stb11* (1BS) | 2 | Chartrain et al. 2005c |
| Salamouni | S | Canada | *StbSm3* (3AS) + *Stb13* (7BL) + *Stb14* (3BS) | 1,2 | Cowling 2006; Cuthbert 2011 |
| Arina<sup>c</sup> | W | Switzerland | *Stb15* (6AS) | 2 | Arraiano et al. 2007 |
| Riband | W | United Kingdom | *Stb15* (6AS) | 1,2 | Arraiano et al. 2007 |
| M3 | S | USA | *Stb16q* (3DL) + *Stb17* (5AL) | 2 | Ghaffary et al. 2012 |
| KM7 | S | The Netherlands | *Stb16q* (3DL) | 2 | Ghaffary et al. 2012 |
| KM20 | S | The Netherlands | *Stb16q* (3DL) + *Stb17* (5AL) | 2 | Ghaffary et al. 2012 |
| KM41 | S | The Netherlands | *Stb17* (5AL) | 2 | Ghaffary et al. 2012 |
| KM73 | S | The Netherlands | *Stb16q* (3DL) + *Stb17* (5AL) | 2 | Ghaffary et al. 2012 |
| Monon | W | USA | Susceptible check | 1 | Shaner and Finney 1982 |
| Arthur 71 | W | USA | Susceptible check | 1 | Van Ginkel 1986 |
| Tyler | W | USA | Susceptible check | 1 | Christ and Frank 1989 |
| Clark | W | USA | Susceptible check | 1 | Adhikari et al. 2004b |
| Patterson | W | USA | Susceptible check | 1 | Adhikari et al. 2004b |
| Taichung 29 | S | Japan | Susceptible check | 1,2 | Kema et al. 2000 |

<sup>a</sup> W = winter type growth; S = spring type growth; F= facultative spring type growth
<sup>b</sup> Experiment 1 = screening of 5 differential wheat lines plus 6 susceptible check cultivars with 13 isolates of *Z. tritici*; Experiment 2 = screening of 21 differential wheat cultivars with the T2 and T46 isolates of *Z. tritici*
<sup>c</sup> These lines also carry the *Stb6* gene
<sup>d</sup> These lines also carry the *Stb7* gene

In the first experiment, seedlings of 11 wheat lines with known and empirical genes for resistance to STB (Table 1) were inoculated with 12 isolates of *Z. tritici* sampled from diverse origins in the U.S. plus the sequenced reference for bread wheat isolates, IPO323, sampled from the Netherlands (Table 2). This assay was conducted to select discriminating isolates for a subsequent seedling test of the soft red winter wheat panel population. Based on the results of these initial inoculations, two isolates of *Z. tritici* collected originally in 1995, T46 (synonym: IN95-Lafayette-1196-WW 1-2) and T2 (synonym: MN95-Marshall-Stephen-SW 3-1), sampled near West Lafayette in Tippecanoe County, Indiana and near Stephen in Marshall Co., Minnesota, respectively, were consequently selected and used to screen seedlings of the soft red winter wheat panel population for STB resistance during experiment two.

**Table 2.** Collection information about the 13 isolates of *Zymoseptoria tritici* used for resistance testing^a^.

| Isolate ID | Synonym | Wheat growth type<br>or cultivar | Collection location |
| --- | --- | --- | --- |
| T2 | MN95-Marshall-Stephen-SW 3-1 | Spring | Marshall Co., MN, USA |
| T4 | MN95- Marshall-Stephen-SW 6-1 | Spring | Marshall Co., MN, USA |
| T5 | MN95- Marshall-Stephen-SW 7-1 | Spring | Marshall Co., MN, USA |
| T24 | ND95-Cavalier-Langdon-SW 6-3 | Spring | Cavalier Co., ND, USA |
| T25 | ND95-Ward-Minot-SW 1-1 | Spring | Ward Co., ND, USA |
| T37 | ND95-Ward-Minot-SW 5-2 | Spring | Ward Co., ND, USA |
| T40 | OH95-Henry-Custar-SRW 2-1 | Winter | Henry Co., OH, USA |
| T42 | OH95-Henry-Custar-SRW 2-3 | Winter | Henry Co., OH, USA |
| T45 | IN95-Lafayette-1196-WW 1-1 | Winter | Tippecanoe Co., IN, USA |
| T46 | IN95-Lafayette-1196-WW 1-2 | Winter | Tippecanoe Co., IN, USA |
| T47 | IN95-Lafayette-1196-WW 1-3 | Winter | Tippecanoe Co., IN, USA |
| T48 | IN95-Lafayette-1196-WW 1-4 | Winter | Tippecanoe Co., IN, USA |
| IPO323 | — | Arminda | The Netherlands |

<sup>a</sup> All cultures were isolated from infected leaves collected during the 1995 growing season except for IPO323, which was obtained in 1981

The third experiment involved inoculating seedlings of 21 differential hexaploid wheat lines containing 17 previously mapped *Stb* genes plus susceptible controls with the T2 and T46 isolates of *Z. tritici* (Table1). Results of this last seedling experiment were used to postulate resistance genes in the soft red winter wheat panel population.

### Experimental design, inoculation procedure and plant maintenance

All seedlings were tested for STB resistance in a completely randomized design in triplicate under the controlled conditions of a growth chamber. The susceptible check cv. Taichung 29 was included in all seedling tests.

Screening tests with the 11 and 21 wheat differential lines were grown in rows in 2.5 x 2.5 x 3.5 cm (length, width, height) Coex square plastic pots (Greenhouse Megastore, USA), whereas the 337 soft red winter wheat panel lines were planted in flats containing 50 pots each (Greenhouse Megastore, USA).

Flats were considered as plots and individual lines were the experimental units. For each isolate/replicate combination, wheat lines were randomly arranged in the flats. Each flat included 48 soft red winter wheat panel lines and the susceptible cultivar Taichung 29 that was in duplicate. The randomization layout was generated using the *Agricolae* package in the R environment (de Mendiburu 2014; Team 2013).

For all assays, five seeds were planted per pot and subsequently thinned to three well-developed, homogeneous seedlings. Seeds were sown in Promix potting soil, a mixture formulated for growing plants indoors (Greenhouse Megastore, USA). Plant development was allowed for 10 days in a growth chamber adjusted to a temperature of 22°C and a relative humidity (RH) of 70%, under a fluorescent light temperature of 6400K and a photoperiod of 16/8 h (light/dark) prior to inoculation.

Inoculum was initiated by pre-cultures of each isolate grown in an autoclaved 100-ml Erlenmeyer flask containing 50 ml of yeast glucose (YG) liquid medium (30 g of glucose, 10 g of yeast per liter of demineralized water). Flasks were inoculated with frozen isolate samples that were taken directly from the *Z. tritici* isolate culture collection maintained by the USDA-Agricultural Research Service on the Purdue University campus, stored at –80°C. Flasks were placed in a Lab-Line orbital shaker set at 120 rpm at room temperature (∼22°C) for 5–7 days. These pre-cultures were then used to inoculate four 1-L Erlenmeyer flasks containing 500 ml of YG medium per isolate that were incubated under the aforementioned conditions to provide sufficient inoculum for the seedling inoculation assays at growth stage (GS) 11 (Zadoks et al. 1974). Spores were collected after overnight settling in static cultures, concentrated by decanting the supernatant medium, adjusted to 1 x 10^7^ spores/ml in a total volume of 40 ml for a set of 18 pots with a haemocytometer and supplemented with two drops of Tween 20 surfactant.

Inoculations were conducted by spraying the spores over the 10-day-old seedlings with a hand-held sprayer. Inoculated plants were incubated in transparent plastic bags for 48 h at 100% RH in the growth chamber. Beginning ten days after inoculation, seedlings were trimmed to remove the second and subsequent leaves to enable sufficient light to reach the inoculated primary leaves for appropriate disease development. An instant soluble fertilizer (Miracle-Gro, USA) (0.5 g/l) was applied to saturation once at 10 days post inoculation to maintain plant condition.

### Data collection and statistical analysis

Disease severities were evaluated at 15, 18 and 22 days post inoculation (dpi) for three leaves per genotype over three replications for 9 leaves in total. These multiple observations enabled Area Under the Disease Progress Curve (AUDPC) calculations for quantitative analyses of temporal differences in disease progress. We estimated the quantitative presence of necrosis (N) and pycnidia (P) on the inoculated seedling leaves visually in percentages. AUDPC calculations for seedling scores followed the trapezoidal method, which approximates the time variable and calculates the average disease intensity between each pair of adjacent time points (Madden et al. 2007).

Analysis of variance was performed in the R environment (Team R 2013) on all generated phenotypic data to determine the effects of line, isolate and any line x isolate interactions. Significance of differences between isolates was determined using the least significant difference (LSD) function in the *Agricolae* package in the R environment (de Mendiburu 2014; Team R 2013). Isolate x line groupings of N and P AUDPC scores were defined based on the Bonferroni test at *p* <0.05. Box-plot figures were generated in the R environment using the boxplot function (Team R 2013).

For the N and P AUDPC scores generated on the 337 soft red winter wheat panel lines, a first check of the data was performed to assess potential variation among the three replicates for each isolate using the lme4 package in the R environment (Bates et al. 2015; Team R 2013). Data variation was checked based on the common susceptible cultivar Taichung 29. Subsequently, a mixed model was generated using the “lmer” command in the lme4 package on all lines over the three replicates. Lines were the fixed effect term of the model and replicates constituted the random-effect term. The least-squares means (LS means) of the N and P AUDPC scores of lines were estimated using the “fixef” statement in lme4, and were used as an estimate of phenotype for subsequent marker-trait association analyses.

Heritability (*h^2^*) was estimated for N and P traits based on the line means as follows:

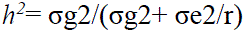

where σg2 is the variance component due to genotypes, σe2 is the variance component due to unexplainable error, and r is the number of replicates, which is three per isolate.

Pearson correlation coefficients were determined for the N and P AUDPC LS mean scores of the soft red winter wheat panel lines inoculated with the T2 and T46 isolates of *Z. tritici* using the “Cor” function and visualized using the “Corrplot” package in the R environment (Team 2013; Wei et al. 2017a). Frequency distribution and the density curve figure were generated using the “hist” function in the R environment (Team 2013).

### Genotyping and population structure

Genomic DNA was extracted from leaves sampled from single plants grown in a greenhouse. Reduced-representation libraries were generated by *Pst1-Msp1* digestion following the protocol of Poland et al. (2012). Samples were pooled in 144-plex multiples to create libraries and each library was sequenced on a single lane of an Illumina Hi-Seq 2500. SNP calling was performed using the Brown5 GBSv2 pipeline1 using 64-base kmer length and minimum kmer count of 5 (Daba et al. 2018; Glaubitz et al. 2014), and reads were aligned to the IWGSC RefSeq assembly v1.0 (Alaux et al. 2018). This resulted in 169,486 unfiltered single-nucleotide polymorphisms (SNPs) that were identified after variant calling. These SNPs were subsequently filtered for missing data (≥20 %) and for minor allele frequency (MAF) of 5% or less. Missing genotypic data were imputed using the Linkage Disequilibrium K-number neighbor imputation method (LDKNNi) algothirm (Money et al. 2015) implemented in TASSELv 5.2 (Bradbury et al. 2007). In total, 14,341 SNPs were retained after filtering for MAF and missing data and were used in this study.

Population structure was evaluated using principal component analysis (PCA) of the 14,341 SNP markers, implemented in TASSEL5.0 (Bradbury et al. 2007). Accordingly, the eigenvalues of all the principal components and the proportions of individual eigenvalues to the total variance (component contribution rates) were calculated. To illustrate relationships among lines, a three-dimensional plot and a kinship matrix were derived from the GAPIT package in the R environment 3.4.3 (Lipka et al. 2012). This analysis was performed using a compressed mixed linear model (CMLM) that replaces the genetic effects of individuals with those of the group to which each individual belongs (Zhang et al. 2010). The marker-based kinship matrix was calculated using the VanRaden method and then used to create a clustering heat map of the soft red winter wheat panel population in the GAPIT R package. Hence, individuals were clustered into groups on the basis of their relationships derived from all the available genetic markers (Tang et al. 2016). The first three PCs were displayed in a three-dimensional plot and the kinship matrix was displayed as a heat map, where red indicates the highest correlation between pairs of individuals and yellow indicates the lowest correlation. A hierarchy tree among individuals was displayed based on their kinship (Tang et al. 2016).

### Genome-Wide Association analysis

Genome-Wide Association (GWA) analysis was computed using the 14,341 SNPs and the least-square means of the N and P AUDPC scores. Two models were implemented in the R environment to identify markers associated with the *Z. tritici* resistance, and results were compared. Consistent marker-trait associations (MTAs) between both models were considered.

At first, a Fixed and random model Circulating Probability Unification (FarmCPU) algorithm was implemented to identify *Z. tritici* resistance MTAs (Liu et al. 2016a). This approach used a Multiple Loci linear Mixed Model (MLMM) that incorporates multiple markers simultaneously as covariates in a stepwise Mixed Linear Model (MLM) to partially remove the confounding between testing markers and kinship (Liu et al. 2016a). Subsequently, a Compressed Mixed Linear Model (CMLM) was implemented in the GAPIT package available in the R environment (Tang et al. 2016). This model clusters the individuals into groups and fits genetic values of groups as random effects in the model, thus reducing false negatives and enhancing the identification of true associations (Zhang et al. 2010).

For both implemented models, the cut-off of significant association was based on a False Discovery Rate (FDR) in the R environment. An MTA was considered significant at *-log (p)* > 4.0. Manhattan plots and the quantile-quantile (QQ) plot figures were subsequently derived from the FarmCPU and CMLM GAPIT output.

### Candidate genes for *Z. tritici* resistance

Linkage disequilibrium (LD) was calculated based on the 14,341 SNP markers using the Tassel 5 software as described by Gaire et al. (2020). LD was quantified using the squared allele frequency correlation (R²). The determination of the LD decay threshold was based on the reported threshold in this soft wheat panel as described by Gaire et al. (2020) to guide the definition of LD blocks in the population. Based on these parameters, LD blocks surrounding significant marker–trait associations (MTAs) were identified to delineate putative genomic regions associated with the traits of interest. Candidate genes for *Z. tritici* resistance were subsequently investigated using the wheat reference genome IWGSC RefSec v1.0 (Alaux et al. 2018).

## Results

### Phenotypic data analysis

The Z*ymoseptoria tritici* isolates grew successfully under laboratory conditions enabling appropriate inoculum production and phenotyping assays. None of the tested cultivars was resistant to the entire suite of *Z. tritici* isolates, resulting in a significant isolate-by-cultivar interaction, indicating specific gene action. Interestingly, necrosis scores were high in most of the tested lines and were less variable compared to pycnidia scores that were highly variable (Supplementary Table 1, Fig. 1). This observation was confirmed by the analysis of variance, indicating a higher significant difference for pycnidia scores compared to those for necrosis (Table 3). The Isolate x Line term was large, emphasising that the observed phenotypic variation was mainly due to the action of specific genes (Table 3).

**Fig 1.**
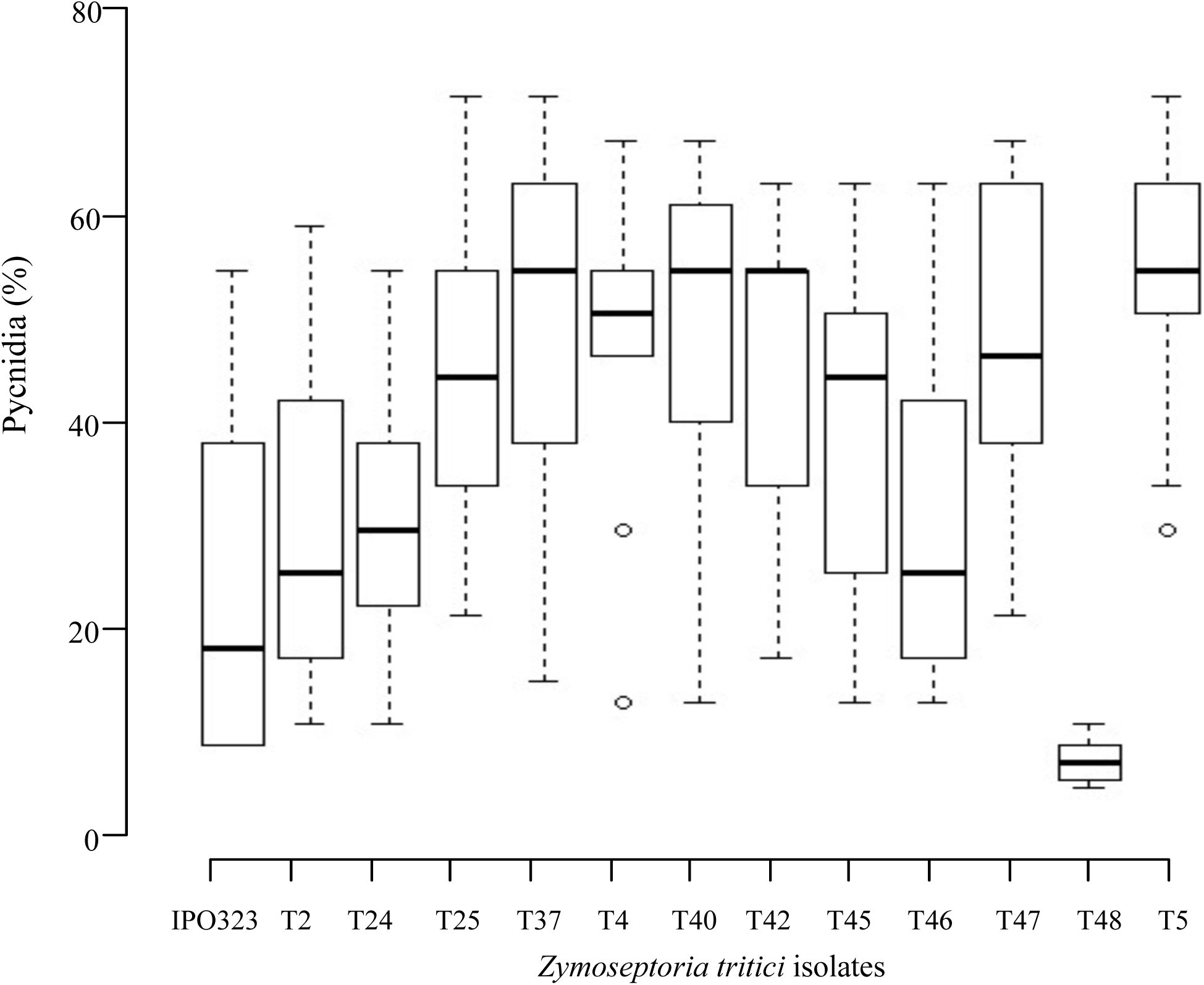
Box plots of percentage pycnidia coverage on resistant and susceptible cultivars of wheat inoculated with 13 diverse isolates of *Zymoseptoria tritici*. Dots outside the main plots indicate outliers.

**Table 3.** Analysis of variance of the necrosis and pycnidia scores on seedlings with known and predicted sources of resistance against Septoria tritici blotch tested with 13 isolates of *Zymoseptoria tritici* sampled from diverse origins.

| Source of variance | Df<sup>a</sup> | Sum of squares |  | Mean Squares |  | F value |  | Pr(>F) <sup>b</sup> |  |
| --- | --- | --- | --- | --- | --- | --- | --- | --- | --- |
|  |  | Necrosis | Pycnidia | Necrosis | Pycnidia | Necrosis | Pycnidia | Necrosis | Pycnidia |
| Isolate | 12 | 1895 | 32247 | 157.9 | 2687.2 | 1.6 | 10.8 | 0.1* | 5.63E-14*** |
| Genotype | 10 | 1660 | 14396 | 166.0 | 1439.6 | 1.7 | 5.8 | 0.1* | 5.72E-07*** |
| Isolate x genotype | 120 | 10886 | 28192 | 96.3 | 249.5 | 1.5 | 7.8 | 0.1* | 1.35E-10*** |
| Residuals | 123 | 12546 | 42589 |  |  |  |  |  |  |

<sup>a</sup> Degrees of freedom
<sup>b</sup> \*\*\* = significant at $p < 0.001$ ; \* is $< 0.01$

Selection of the *Z. tritici* isolates for subsequent screening of the soft red winter wheat panel lines was based mainly on isolate indicators that might potentially identify previously known *Stb* genes as well as potentially new genes for resistance. Hence, based on the pre-screening results, uniformly virulent or avirulent isolates were not considered further as they had lower ability to discriminate resistance genes (Supplementary Table 1). Based on these criteria, the T2 and T46 isolates of *Z. tritici* were chosen for further seedling screening assays.

The soft red winter wheat panel lines were tested with the T2 and the T46 isolates that were sampled originally from Minnesota and Indiana, respectively. The seedling screening of the soft red winter wheat panel resulted in approximately normal frequency distributions for the necrosis and pycnidia AUDPC scores for both isolates (Fig. 2). A skewed distribution towards a resistant phenotype was observed for pycnidia AUDPC scores for both isolates. In contrast, necrosis scores were higher and skewed towards a susceptible phenotype (Fig. 2).

**Fig 2.**
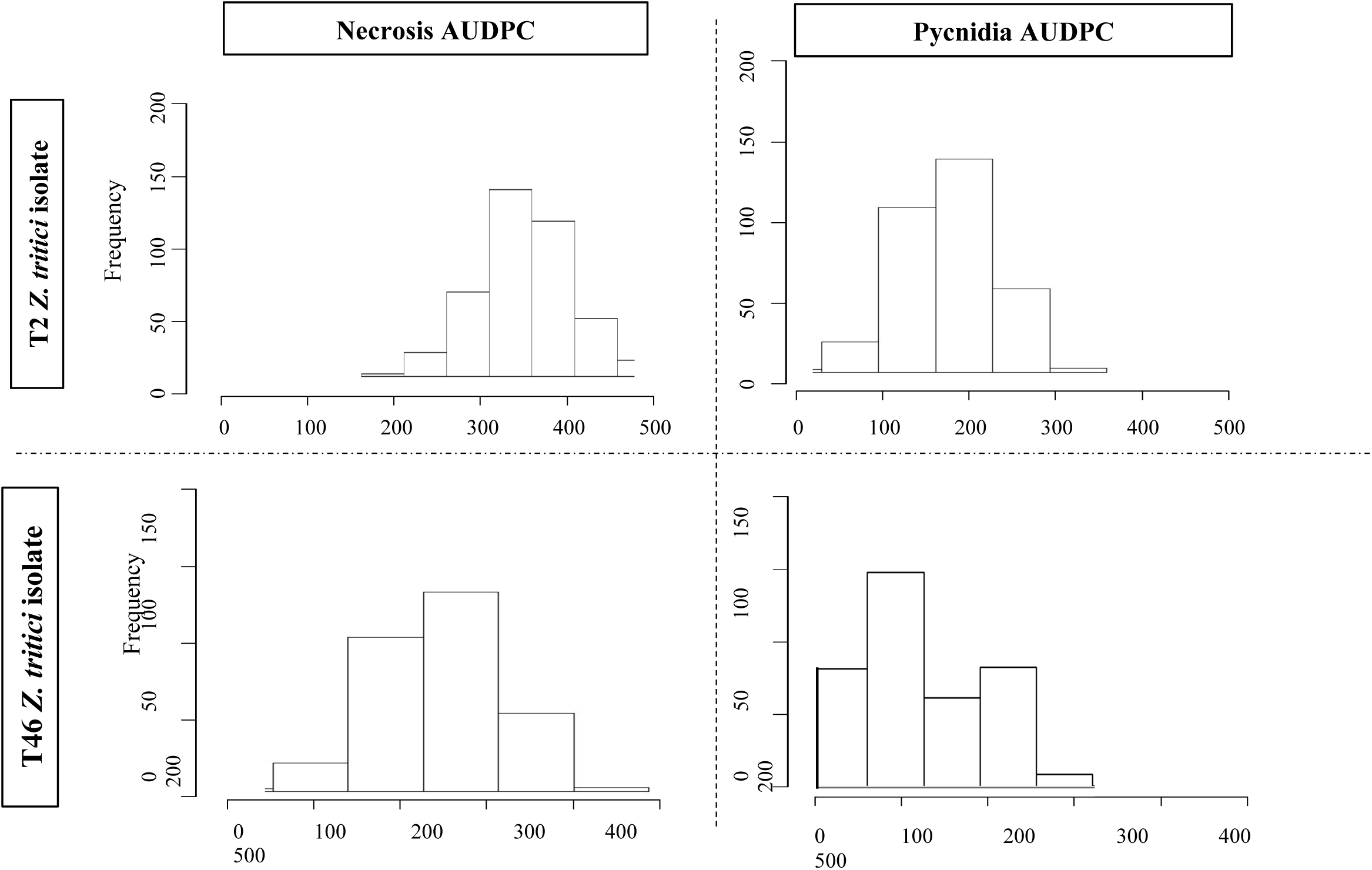
Frequency distribution of the least-square means of the Necrosis and Pycnidia AUDPC scores for the T2 and T46 isolates of *Zymoseptoria tritici* following inoculations of the tested “Purdue Panel” lines under controlled conditions.

Although a similar phenotypic pattern was observed for both isolates, the analysis of variance of the tested soft red winter wheat panel lines indicated that the isolate x genotype term was highly significant, indicating a difference between the tested isolates and an ability to discriminate resistance genes that are specific for each isolate (Table 4). Interestingly and in contrast with the pre-screening results, both necrosis and pycnidia AUDPC scores were highly significant for the isolate x line term (Table 4).

**Table 4.** Analysis of variance of the necrosis and pycnidia Area Under the Disease Progress scores registered on seedlings of the soft red winter wheat panel population inoculated with the T2 and T46 isolates of *Zymoseptoria tritici*.

| Source of variance | Df<sup>a</sup> | Sum of squares |  | Mean Squares |  | F value |  | Pr (>F)<sup>b</sup> |  |
| --- | --- | --- | --- | --- | --- | --- | --- | --- | --- |
|  |  | Necrosis | Pycnidia | Necrosis | Pycnidia | Necrosis | Pycnidia | Necrosis | Pycnidia |
| Isolate | 1 | 1452142 | 112113 | 1452142 | 112113 | 224.1 | 46.6 | <2e-16*** | 4.14e-11*** |
| Genotype | 336 | 2595303 | 815106 | 7724 | 2426 | 1.2 | 1.0 | 0.054 | 0.472 |
| Isolate x genotype | 336 | 2177260 | 808908 | 6480 | 2407 | 3.9 | 3.9 | 1.09e-40*** | 2.15e-11*** |
| Residuals | 673 | 6224705 | 1736126 |  |  |  |  |  |  |

<sup>a</sup> Degrees of freedom
<sup>b</sup> \*\*\* = significant at $p < 0.001$

Differentiation between the isolates was clear in the overall population means and ranges for the necrosis and pycnidia AUDPC scores (Table 5). Overall, higher necrosis and pycnidia AUDPC population means were observed for lines inoculated with the T2 *Z. tritici* isolate compared to those with T46 (Table 5, Supplementary Fig. 1). The soft red winter wheat panel population means of 417.4 for necrosis and 111.1 for pycnidia AUDPC scores for the T2 isolate were higher compared to the values of 324.5 and 85.3 for the same statistics on lines inoculated with the T46 isolate. A high variability demonstrated by large necrosis and pycnidia AUDPC ranges also was observed between lines inoculated with the same isolate (Table 5). However, both isolates gave the same low and moderate heritabilities for necrosis and pycnidia AUDPC scores of 0.2 and 0.4, respectively (Table 5).

**Table 5.** Summary of the necrosis and pycnidia development scores plus their heritabilities when scored on the soft red winter wheat panel population inoculated with the T46 and T2 isolates of *Zymoseptoria tritici* at the seedling stage.

| Parameter | $h^2$<sup>a</sup> | | Population mean | | Range | | | |
| --- | --- | --- | --- | --- | --- | --- | --- | --- |
|  |  |  |  |  | Minimum |  | Maximum |  |
|  | T46 | T2 | T46 | T2 | T46 | T2 | T46 | T2 |
| Necrosis AUDPC<sup>b</sup> | 0.2 | 0.2 | 324.5 | 417.4 | 88.3 | 25 | 520 | 600 |
| Pycnidia AUDPC<sup>b</sup> | 0.4 | 0.4 | 85.3 | 111.1 | 16.7 | 10 | 283.3 | 255 |

<sup>a</sup> Heritability
<sup>b</sup> Area Under the Disease Progress Curve

A different correlation pattern was observed between the isolates for the two parameters; necrosis and pycnidia AUDPC scores were highly correlated for the T2 isolate (0.69), but were only correlated weakly for the T46 isolate (0.27) (Fig. 3). Weak correlations also were observed between the same disease parameters (necrosis or pycnidia) and the different isolates (Fig. 3).

**Fig 3.**
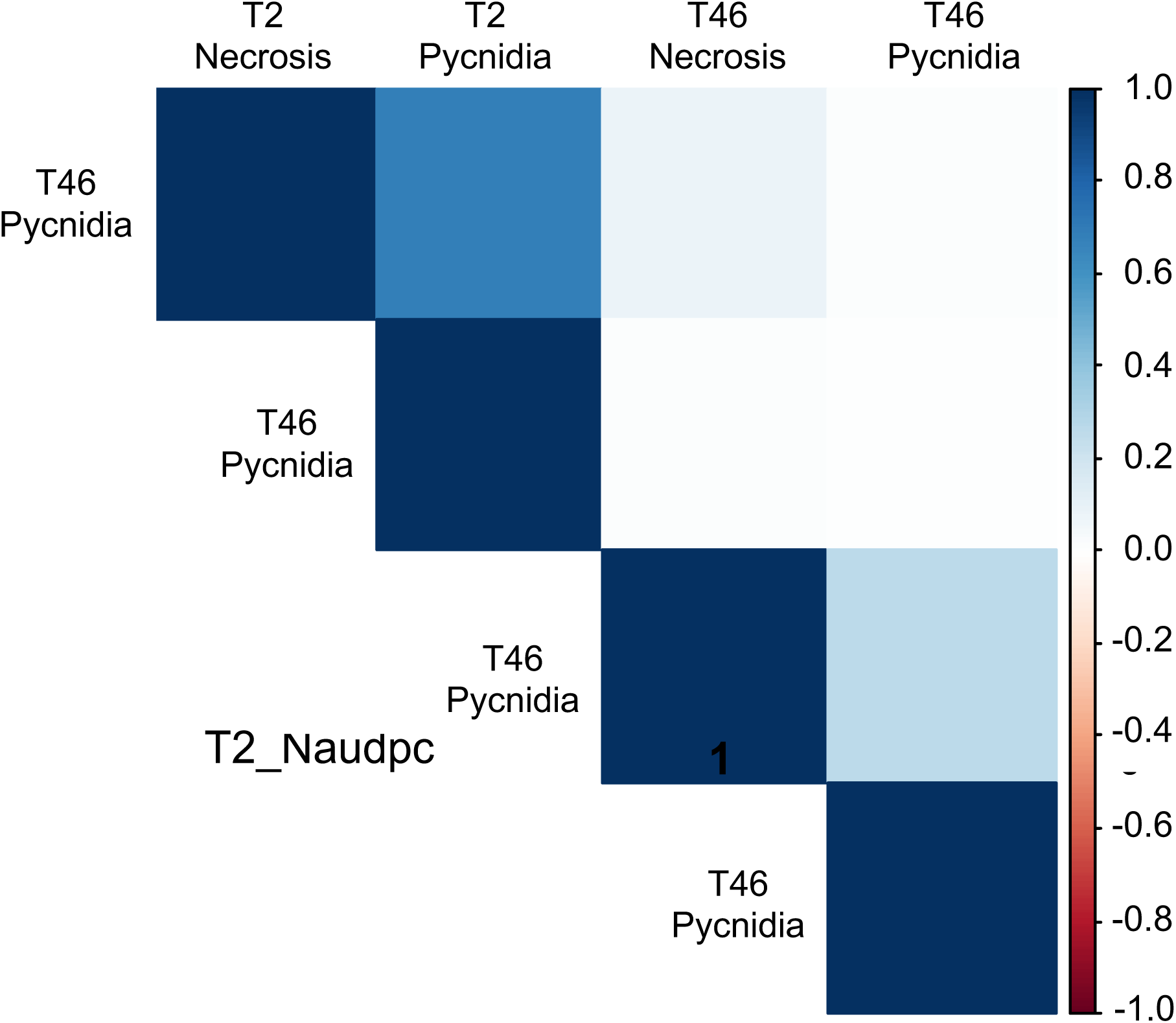
Pearson correlation coefficients for Area Under the Disease Progress Curve (AUDPC) between the indicated phenotypic traits after inoculation with two isolates of *Zymoseptoria tritici*. Each trait is named by the isolate ID over Necrosis or Pycnidia depending on what was scored. A scale bar showing the strength and direction of the correlations is on the right with positive correlations in blue, negative correlations in red and no correlation in white. The dark blue diagonal squares were each trait by itself so were always equal to 1

### Population structure

To evaluate population structure, principal component analysis (PCA) was performed in TASSEL 5 using 14,341 SNP markers, and a kinship matrix was generated in the GAPIT package. Based on model complexity, the panel of 337 wheat lines was classified into four subpopulations (Fig. 4). A marked change was observed at the fourth principal component, which corresponded to a cumulative variance contribution of 25.7% (Fig. 4A).

**Fig 4.**
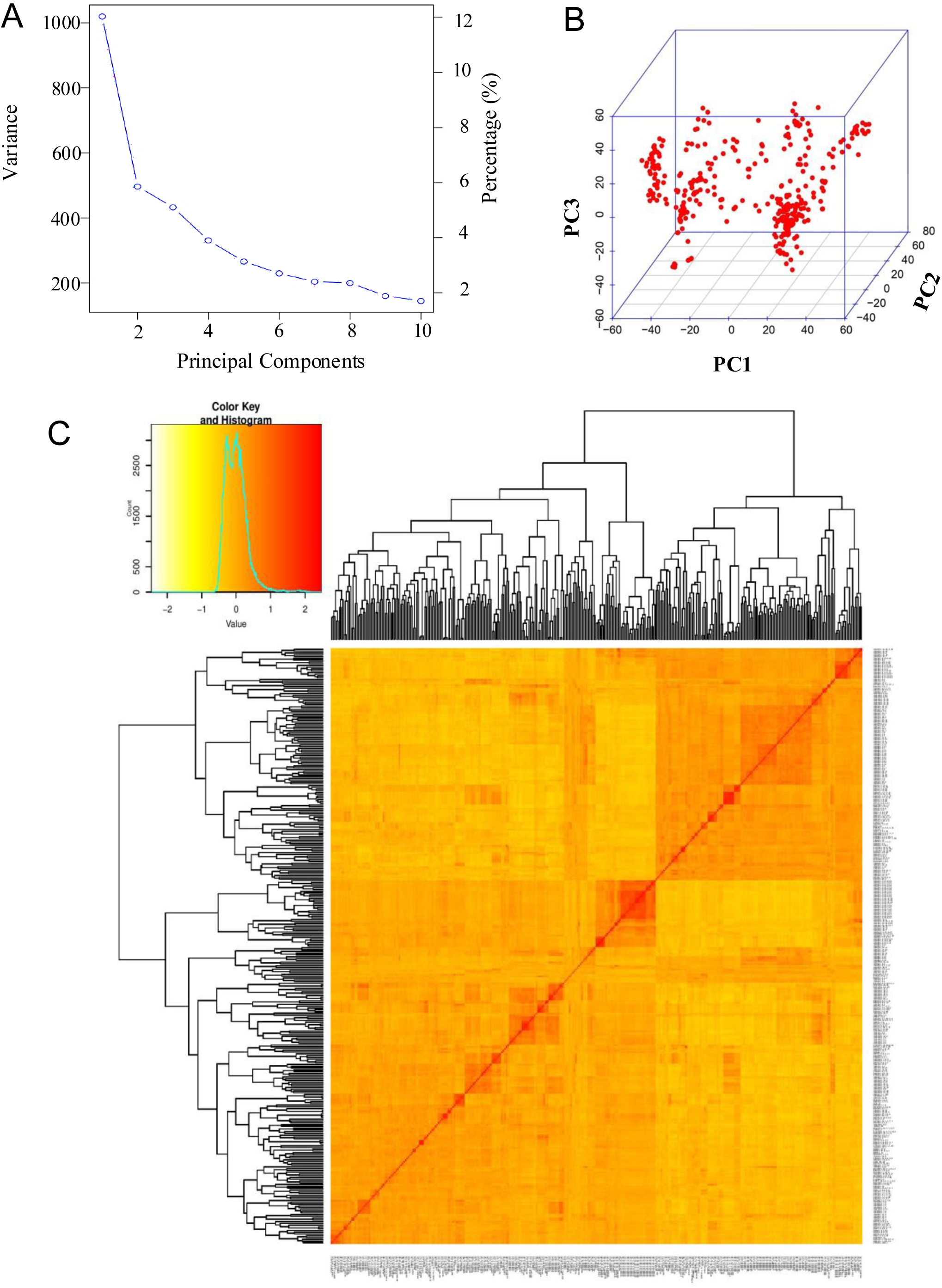
Population structure results based on a principal component analysis (PCA) of the 337 soft red winter wheat panel lines and a heatmap and dendrogram of the kinship matrix estimated using 14,341 Single-Nucleotide Polymorphism (SNP) markers. **A**, changes in variances in each principal component. **B**, a three-dimensional plot of the first three principal components. **C**, kinship analysis with the genetic clustering heatmap created with a matrix for evaluating the genetic differences among the 337 soft red winter wheat panel lines. The red color indicates the highest correlation between pairs of individuals and yellow indicates the lowest correlation. A hierarchy tree among individuals was displayed based on their kinship

The first four principal components together explained 25.7% of the total genetic variation, with PC1 accounting for 11.8%, followed by PC2 (5.2%), PC3 (5.0%), and PC4 (3.7%) (Fig. 4A). A three-dimensional scatterplot based on the first three principal components further separated the wheat lines into discrete genetic clusters (Fig. 4B).

In contrast, clustering based on the kinship matrix suggested that the soft red winter wheat panel was primarily divided into two major groups. Considerable genetic divergence among lines was evident from the color gradient ranging from red to yellow, representing high to low genetic relatedness among lines, respectively (Fig. 4C).

### Marker-trait associations and *Stb* gene postulations in the soft red winter wheat panel population

Two models were compared to assess the association of SNP markers to *Z. tritici* resistance: FarmCPU, which uses a Multiple Loci linear Mixed Model (MLMM); and the Compressed Mixed Linear Model (CMLM) computed in GAPIT.

Overall, 12 SNP markers mapped to chromosomes 1A, 4A, 6A, 7A, 3B, 5B and 1D were associated with the necrosis and pycnidia resistance against *Z. tritici* (Table 6). All markers that were significantly associated with the *Z. tritici* resistance by the FarmCPU model were confirmed when the phenotypic scores also were analyzed with the CMLM model in GAPIT (Table 6, Fig. 5, Supplementary Figure 2). However, significance levels of the associations were higher for the FarmCPU model compared to CMLM as revealed by the *-log(p)* values (Table 6).

**Fig 5.**
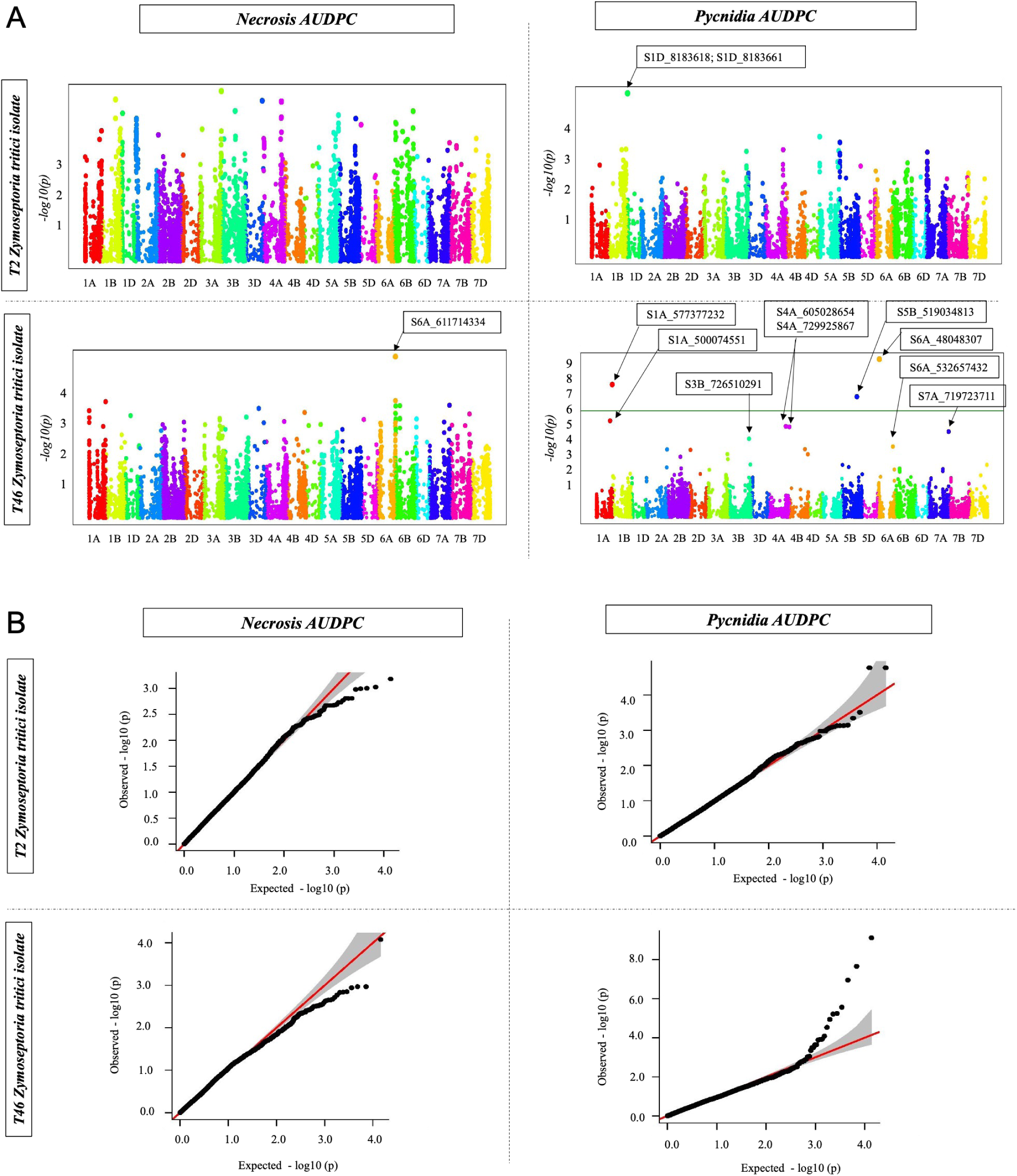
Manhattan plots (panel **A**) and Q-Q plots (panel **B**) generated by the FarmCPU Genome-Wide Association Study analysis. Marker traits associated with necrosis and pycnidia resistance to the T2 and T46 isolates of *Zymoseptoria tritici* in **A** are indicated by arrows. Boxes in **A** include the marker IDs of all Single-Nucleotide Polymorphisms (SNPs) associated with resistance

**Table 6.**
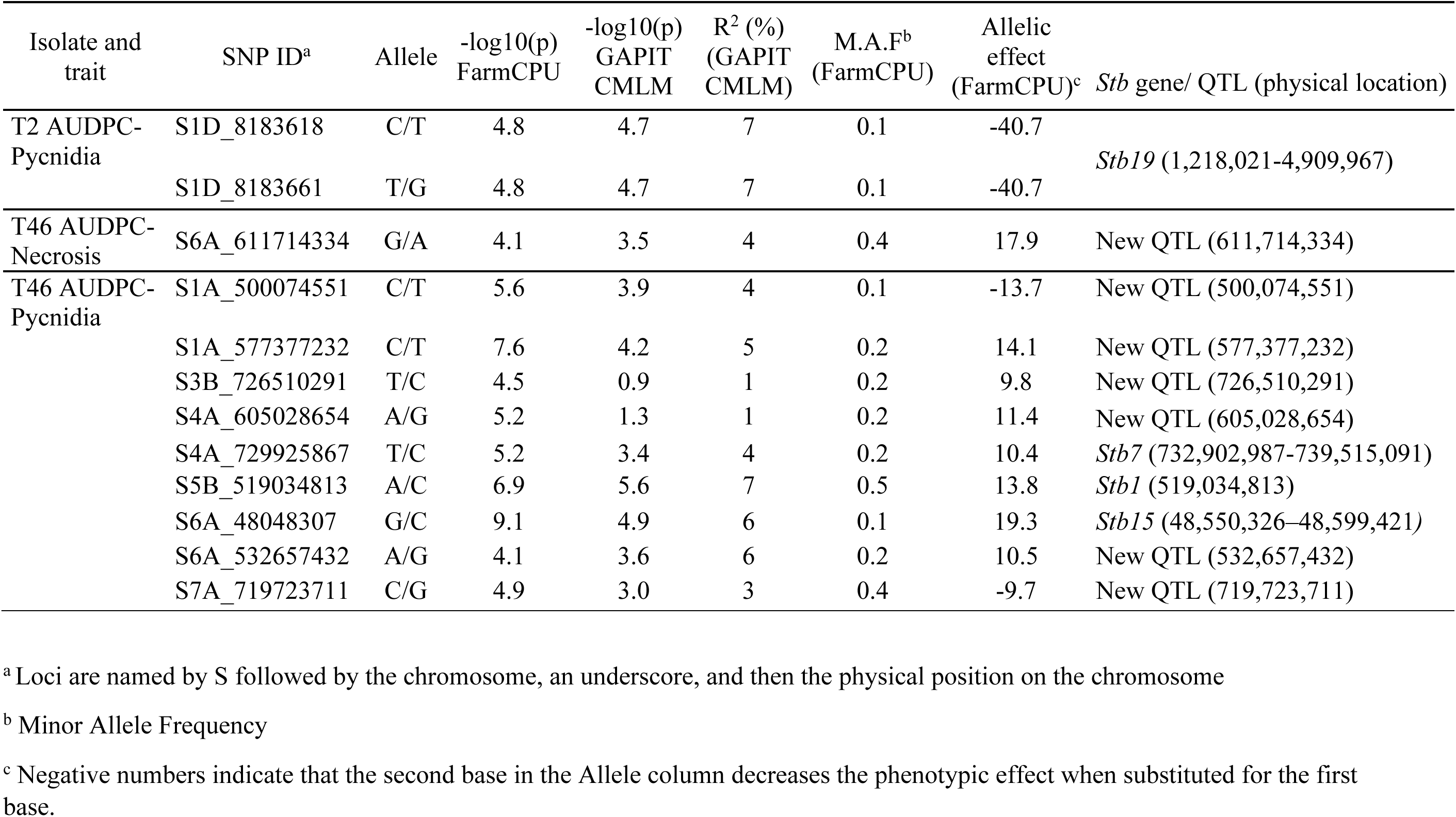
Marker-trait associations with STB resistance mapped on the soft red winter wheat panel population with the T2 and T46 isolates of *Zymoseptoria tritici* at the seedling stage in comparison with the positions of known *Stb* genes and using models from FarmCPU and GAPIT (CMLM)

| Isolate and trait | SNP ID<sup>a</sup> | Allele | -log<sub>10</sub>(p)<br>FarmCPU | -log<sub>10</sub>(p)<br>GAPIT<br>CMLM | R<sup>2</sup> (%)<br>(GAPIT<br>CMLM) | M.A.F<sup>b</sup><br>(FarmCPU) | Allelic<br>effect<br>(FarmCPU)<sup>c</sup> | *Stb* gene/ QTL (physical location) |
| --- | --- | --- | --- | --- | --- | --- | --- | --- |
| T2 AUDPC-Pycnidia | S1D_8183618 | C/T | 4.8 | 4.7 | 7 | 0.1 | -40.7 | *Stb19* (1,218,021-4,909,967) |
|  | S1D_8183661 | T/G | 4.8 | 4.7 | 7 | 0.1 | -40.7 |  |
| T46 AUDPC-Necrosis | S6A_611714334 | G/A | 4.1 | 3.5 | 4 | 0.4 | 17.9 | New QTL (611,714,334) |
| T46 AUDPC-Pycnidia | S1A_500074551 | C/T | 5.6 | 3.9 | 4 | 0.1 | -13.7 | New QTL (500,074,551) |
|  | S1A_577377232 | C/T | 7.6 | 4.2 | 5 | 0.2 | 14.1 | New QTL (577,377,232) |
|  | S3B_726510291 | T/C | 4.5 | 0.9 | 1 | 0.2 | 9.8 | New QTL (726,510,291) |
|  | S4A_605028654 | A/G | 5.2 | 1.3 | 1 | 0.2 | 11.4 | New QTL (605,028,654) |
|  | S4A_729925867 | T/C | 5.2 | 3.4 | 4 | 0.2 | 10.4 | *Stb7* (732,902,987-739,515,091) |
|  | S5B_519034813 | A/C | 6.9 | 5.6 | 7 | 0.5 | 13.8 | *Stb1* (519,034,813) |
|  | S6A_48048307 | G/C | 9.1 | 4.9 | 6 | 0.1 | 19.3 | *Stb15* (48,550,326-48,599,421) |
|  | S6A_532657432 | A/G | 4.1 | 3.6 | 6 | 0.2 | 10.5 | New QTL (532,657,432) |
|  | S7A_719723711 | C/G | 4.9 | 3.0 | 3 | 0.4 | -9.7 | New QTL (719,723,711) |

<sup>a</sup> Loci are named by S followed by the chromosome, an underscore, and then the physical position on the chromosome
<sup>b</sup> Minor Allele Frequency
<sup>c</sup> Negative numbers indicate that the second base in the Allele column decreases the phenotypic effect when substituted for the first base.

No marker-trait associations were mapped in common between the two tested isolates of *Z. tritici* (T2 and T46), highlighting the isolate-specific resistance in the *Z. tritici-*wheat pathosystem (Table 6, Fig. 5, Supplementary Figure 2).

For the T2 isolate of *Z. tritici*, significant marker-trait associations were solely with pycnidia resistance, in contrast to the T46 isolate, which identified markers associated with both necrosis and pycnidia resistance. Distinct markers were associated with necrosis and pycnidia resistances, which confirms that the necrosis and pycnidia phenotypes are likely under distinct genetic control (Table 6, Fig. 5, Supplementary Figure 2).

Two significant SNP markers (*S1D_8183618* and *S1D_8183661*) (-*log (p)*≥ 4.0) that fell into the same LD block were mapped to the short arm of chromosome 1D and were associated with the pycnidia resistance against isolate T2, explaining 7% (R^2^) of the observed phenotypic variance (Table 6, Fig. 5, Supplementary Figure 2). This genomic location overlaps that of the known resistance gene *Stb19*, which was identified in the Australian soft wheat cultivar Lorikeet (Yang et al. 2018).

For the T46 isolate of *Z. tritici*, several chromosomes were associated with the necrosis or pycnidia resistance. Chromosome 6A was associated with both necrosis and pycnidia resistance but with different significant SNP markers. SNP locus *S6A_611714334* was associated with the necrosis resistance and explained 4% of the observed phenotypic variation (Table 6, Fig. 5 and Supplementary Figure 2). Two SNP markers (*S6A_48048307* and *S6A_532657432*) were highly significant for pycnidia resistance and explained 6% of the observed phenotypic variance.

Chromosomes 1A, 3B, 4A, 5B and 7A were solely associated with pycnidia resistance against the T46 isolate. Genomic regions for pycnidia resistance on chromosomes 1A, 3B, 4A and 7A, associated with the SNP markers *S1A_500074551*, *S1A_577377232*, *S3B_726510291*, *S4A_605028654* and *S7A_719723711*, respectively, do not contain any previously mapped *Stb* genes so appear to be novel. The significant markers *S1A_500074551* and *S1A_577377232*, mapped on chromosome 1A, fell into two distinct LD blocks, thus constituting different genomic positions associated with the pycnidia resistance to *Z. tritici*, explaining 4 and 5% of the observed phenotypic variance, respectively. The *S3B_726510291*, *S4A_605028654* and *S7A_719723711* SNP loci on chromosomes 3B, 4A and 7A explained the lowest phenotypic variance with only 1, 1 and 3%, respectively.

Significant SNP loci mapped on chromosomes 4A, 6A and 5B are in genomic regions that overlap with the known *Stb* genes *Stb7*, *Stb15* and *Stb1*, respectively. Two significant SNP markers (*S4A_605028654* and *S4A_729925867*) that were identified on chromosome 4A fell into distinct LD blocks. The MTA identified at 729,925,867 Mbp co-segregated with the known gene *Stb7* identified in the hexaploid wheat line Estanzuela Federal (Brown et al. 2015b; McCartney et al. 2003), explaining up to 4% of the observed phenotypic variance. The other MTA on chromosome 4A may represent a new QTL but only explained 1% of the variance so could be an artifact.

The SNP locus *S6A_48048307* is located approximately 0.50 Mb from the recently cloned *Stb15* locus and falls within the LD decay distance for chromosome 6A (2.73 Mb), suggesting that this association may correspond to the *Stb15* region (Hafeez et al. 2025). In contrast, *S6A_532657432* is located approximately 484.11 Mb from *Stb15*, beyond the LD decay distance, indicating that this MTA is unlikely to be linked to *Stb15* and may therefore represent a distinct, potentially novel QTL associated with Septoria tritici blotch resistance.

Finally, the SNP locus *S5B_519034813*, mapped to chromosome 5B, co-localized with the known resistance gene *Stb1*. MTAs identified on chromosomes 6A and 5B explained 6 and 7% of the observed phenotypic variance, respectively. The *Stb15* and the *Stb1* resistance genes have been reported previously in the wheat lines Arina and Bulgaria 88, identified in Switzerland and the US, respectively (Adhikari et al. 2004a; Arraiano et al. 2007; Czembor et al. 2011). *Stb15* was recently cloned and shown to encode an intronless G-type lectin receptor-like kinase (RLK) (Hafeez et al. 2025).

To confirm the co-segregation of the mapped MTAs on chromosomes 4A, 6A and 5B to the known *Stb* genes *Stb7*, *Stb15* and *Stb1*, a seedling assay was carried out with the T2 and the T46 isolates of *Z. tritici* on a set of wheat differential lines, where a significant isolate × line interaction was observed (Supplementary Table 2, Fig. 6). According to the least significant difference calculated for the necrosis and the pycnidia scores, isolate × line interactions were clustered into distinct groups, which confirmed that the T2 and T46 isolates carry distinct avirulence genes and are potential indicators of known *Stb* genes (Fig. 6). A low pycnidia level was generated by the T46 isolate on cultivars Oasis, Estanzuela Federal and Arina, carrying *Stb1*, *Stb7* and *Stb15*, respectively, which indicates that this isolate is most likely avirulent to those genes. This indicates that the co-localized MTAs on chromosomes 4A, 6A and 5B are likely due to the known genes *Stb1*, *Stb7* and *Stb15*.

**Fig 6.**
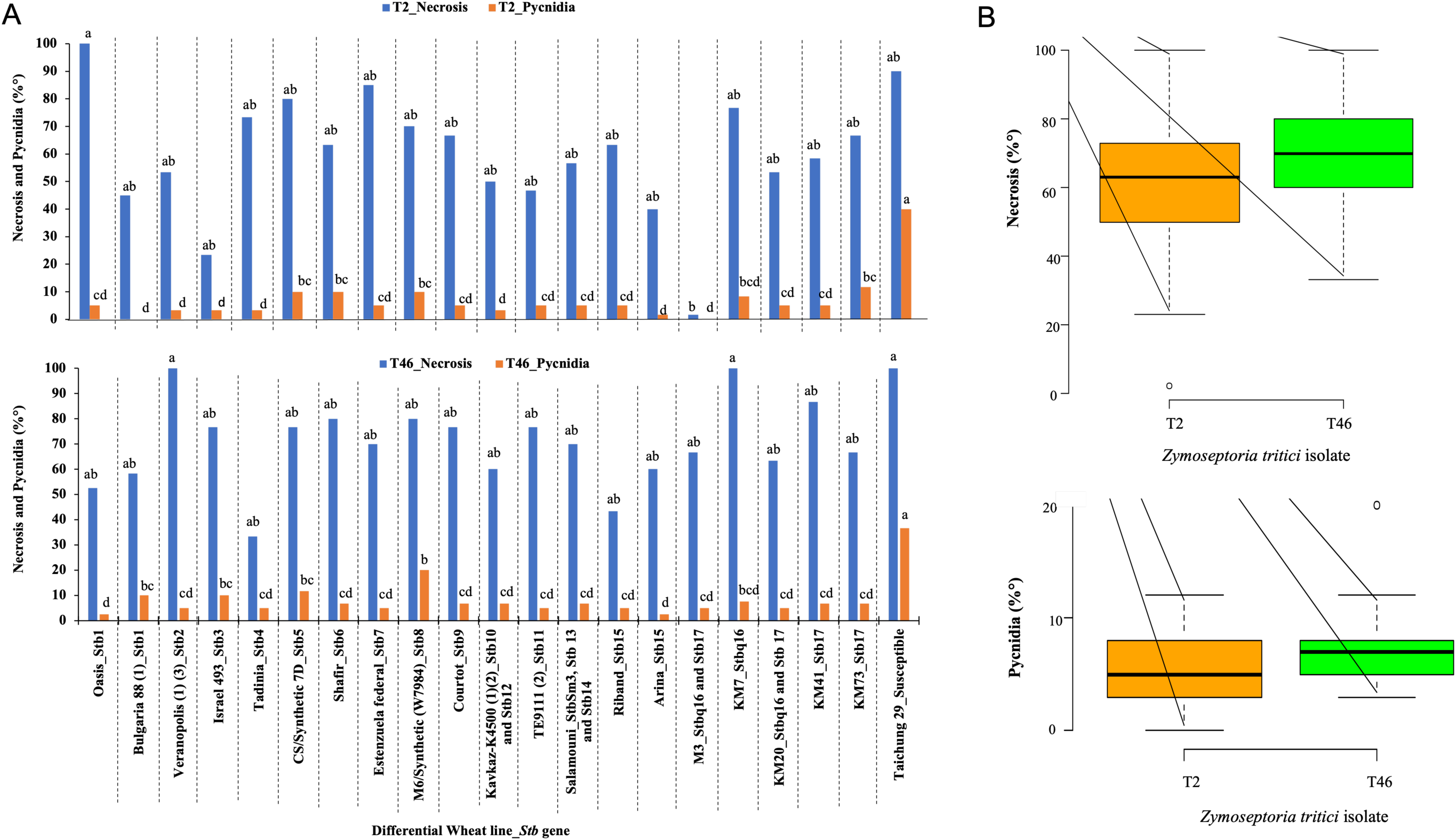
Necrosis and pycnidia percentages that developed on the differential wheat lines inoculated with the *Zymoseptoria tritici* isolates T2 (top panel) and T46 (bottom). **A**, different groups are marked by letters on top of the histogram bars based on the least-square means. **B**, box plots showing differences in necrosis and pycnidia development on the differential wheat cultivars between the T2 and the T46 isolates of *Z. tritici*

### Single time point versus AUDPC comparisons

The single time point analysis sometimes identified additional and different genes compared to those revealed by the AUDPC values calculated over time. Although isolate-specific MTAs were revealed when analyzing both datasets with the two statistical models, a higher number of MTAs was detected when competing necrosis and pycnidia single time-point scores compared to the necrosis and pycnidia AUDPC scores (Supplementary Table 3). Necrosis and pycnidia AUDPC data revealed 12 MTAs associated with the *Z. tritici* resistance, but only two were identified using pycnidia AUDPC scores from the T2 isolate while ten MTAs were associated with necrosis and pycnidia AUDPC resistance against the T46 isolate. In contrast, 14 MTAs were associated with the pycnidia and necrosis scores at a single time point with isolate T2, and 24 MTAs were associated with the T46 isolate when necrosis and pycnidia scores were evaluated at a single time point (Supplementary Table 3).

The accuracy and robustness of both phenotypic datasets to detect markers associated with the *Z. tritici* resistance against isolate T46 were confirmed when using both phenotypic datasets (AUDPC and single time-point scores) (Supplementary Table 3). Nonetheless, the T46 single-time-point phenotypic scores compared in a GWA analysis revealed additional genomic regions, other than those revealed by AUDPC scores, associated with the *Z. tritici* resistance on chromosomes 3A (*S3A_700960980*; *S3A_715967971*), 4B (*S4B_548241937*), 2D (*S2D_29753796*) and 4D (*S4D_439658613*). All the additional MTAs were associated with pycnidial development at 16 (*S4B_548241937*) and 22 dpi (*S3A_700960980, S3A_715967971, S2D_29753796, S4D_439658613*). The additional detected MTAs might be mainly explained by the distinct phenotypic variations when using AUDPC and single-time-point scores generated by the T46 *Z. tritici* isolate.

In contrast to the T46 *Z. tritici* isolate, none of the detected MTAs by the single-time-point scores generated by the T2 isolate of *Z. tritici* were detected by the AUDPC scores. Hence, analyzing the phenotypic variation on different scoring days versus combined into AUDPC led to different GWA results.

### Candidate resistance genes

To provide an initial insight into the types of genes present within the identified MTAs, a preliminary candidate gene search was conducted. Candidate resistance genes were identified within the genomic regions associated with *Z. tritici* resistance based on the linkage disequilibrium of each chromosome (Gaire et al. 2020). Among all of the candidate genes, many fell into three predominant categories of possible resistance genes (Fig. 7, Table 7, Supplementary Tables 4 and 5, Supplementary Figure 3). Predominant candidate resistance gene categories were as follows:

(i) Disease resistance candidates that comprised the leucine-rich-repeat (LRR) and nucleotide-binding site–leucine-rich repeat (NBS-LRR) protein families. This category represents 28% of the putative identified resistance genes and comprised the highest number of candidate resistance genes on chromosomes 4A, 6A (20862967-75370938) and 1D (Fig. 7, Table 7).
(ii) The kinase/kinesin proteins that represent 19% of the identified putative resistance genes. This category is highly represented on chromosomes 6A (252830745-617935868) and 5B (Fig. 7, Table 7).
(iii) Receptor lectin kinases constituted 14% of the total putative resistance genes and were the most common candidate resistance genes on chromosomes 7A and 3B (Fig. 7, Table 7).

**Fig 7.**
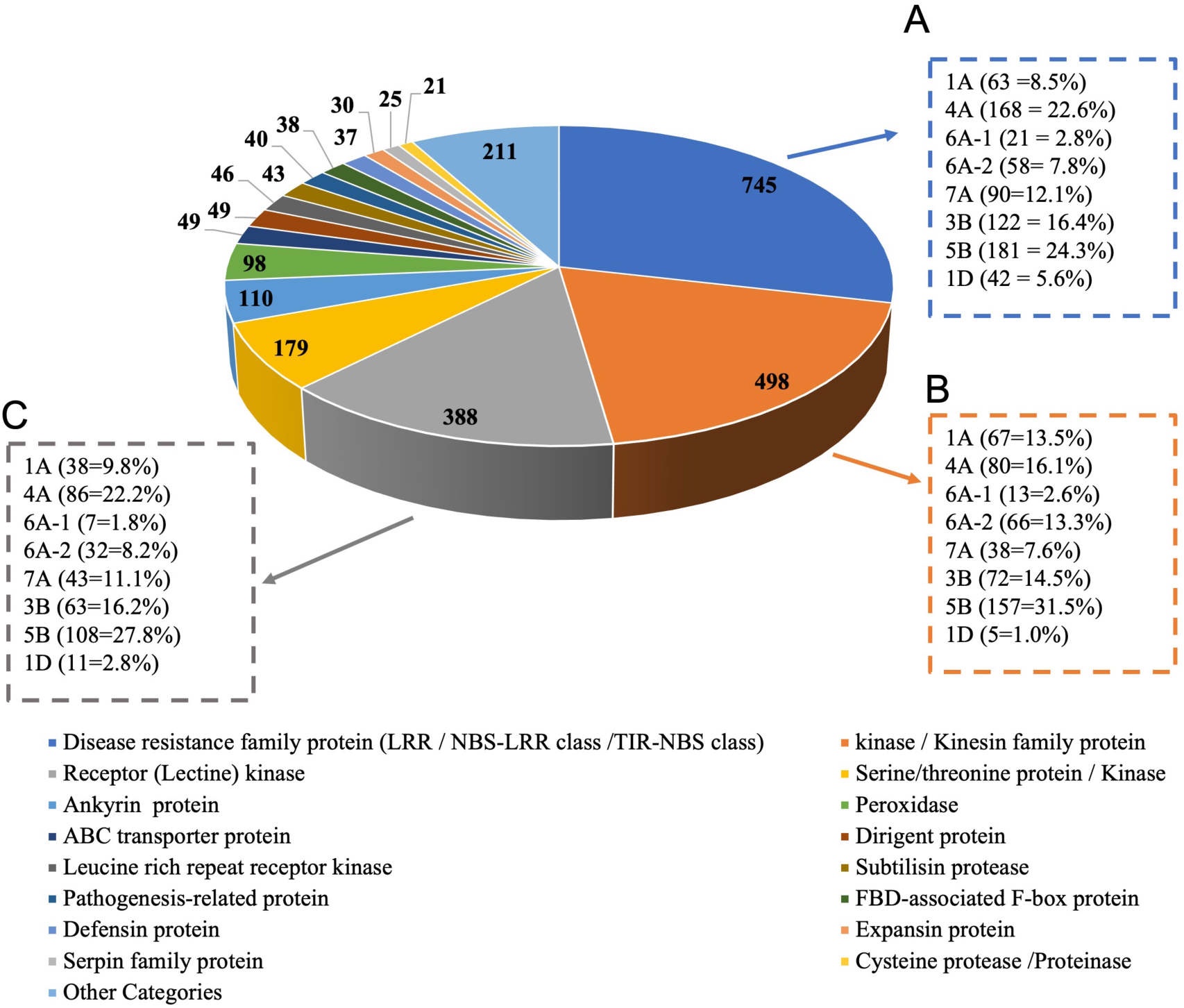
Categories for candidate genes identified in the genomic regions associated with resistance against *Zymoseptoria tritici* from the Genome-Wide Association analysis. Numbers and percentages of the predominant categories per chromosomal region associated with the *Z. tritici* resistance are indicated in panels **A**, **B** and **C**, which represent the disease-resistance protein families, the kinase/kinesin proteins and the receptor lectin kinases, respectively

**Table 7.**
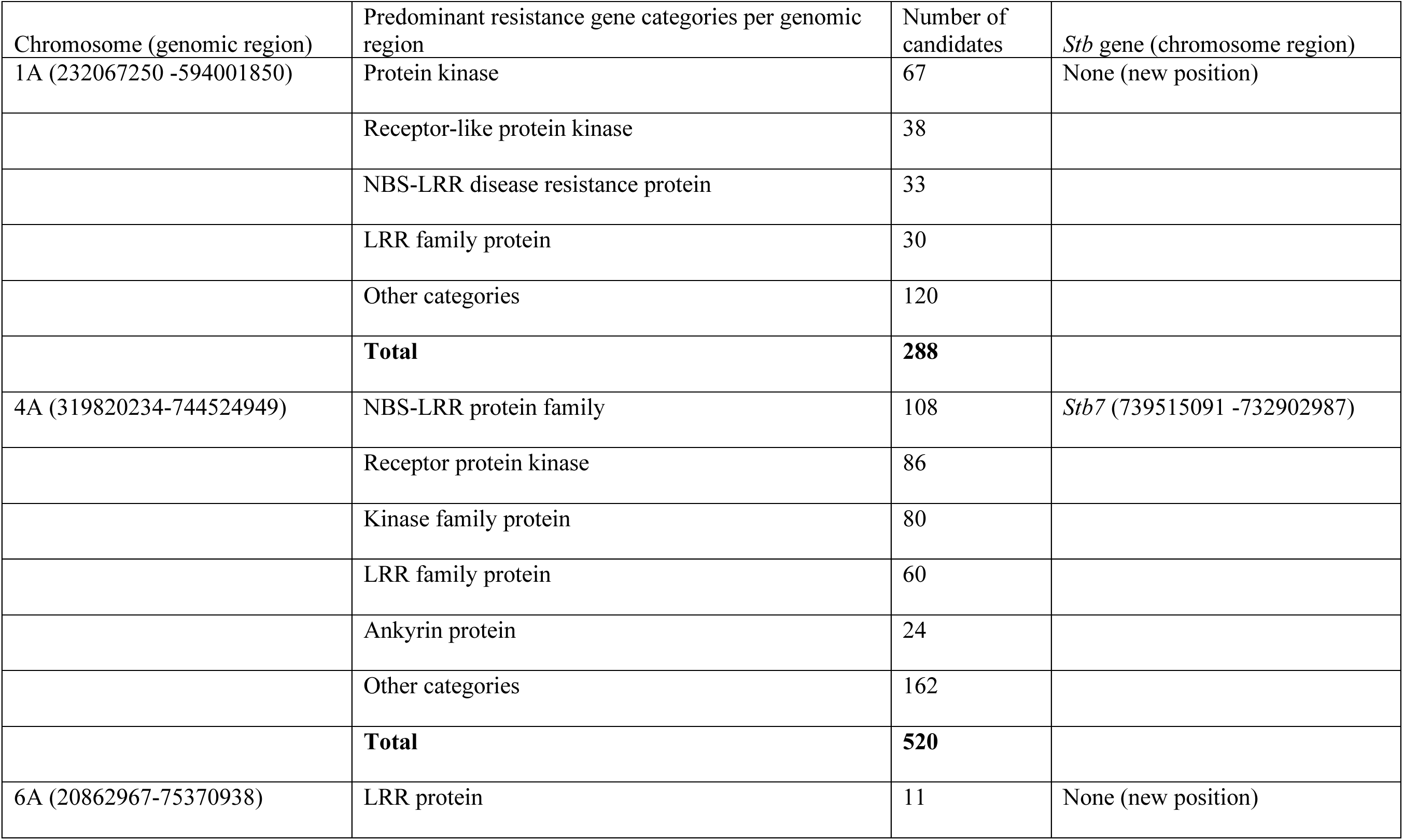
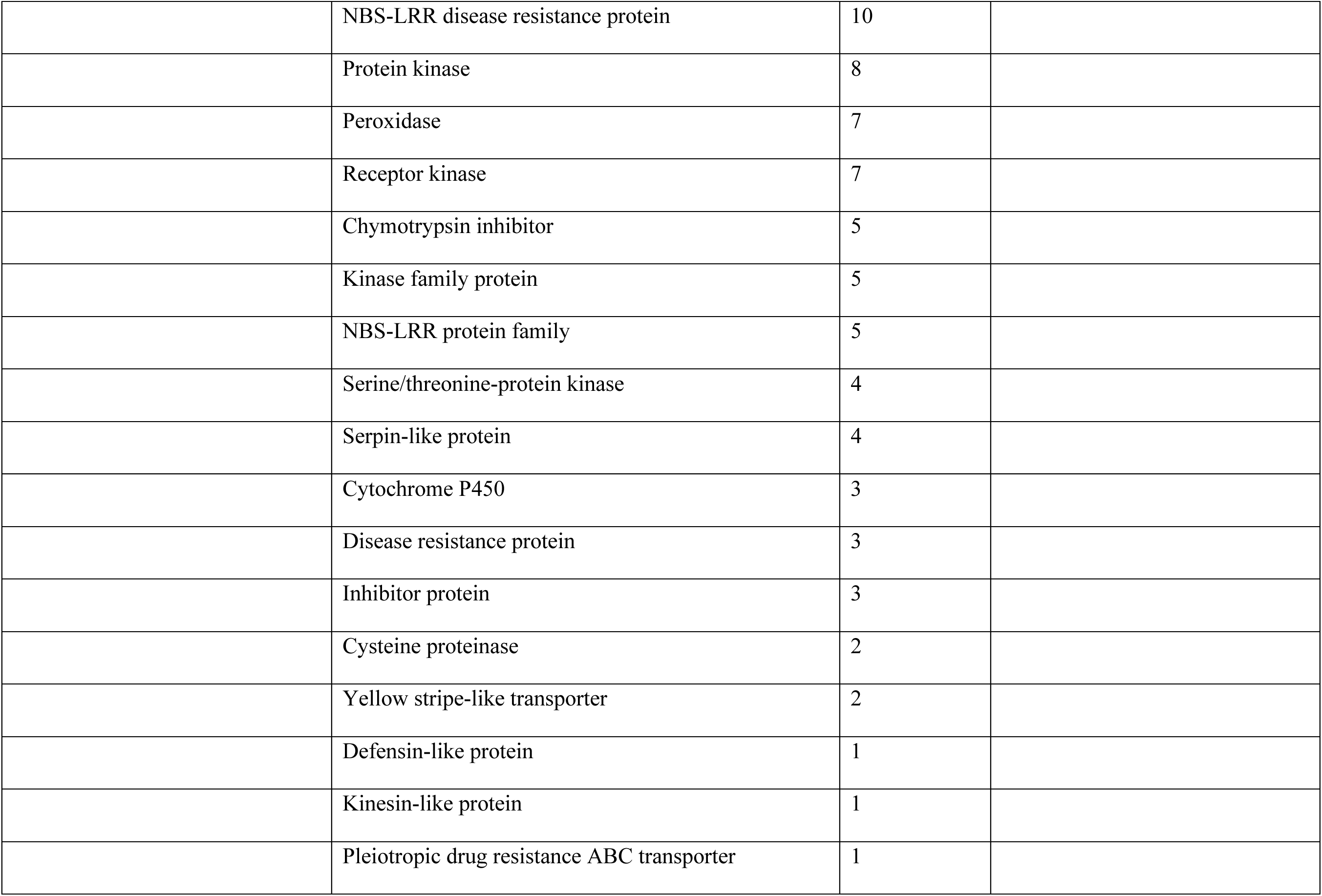
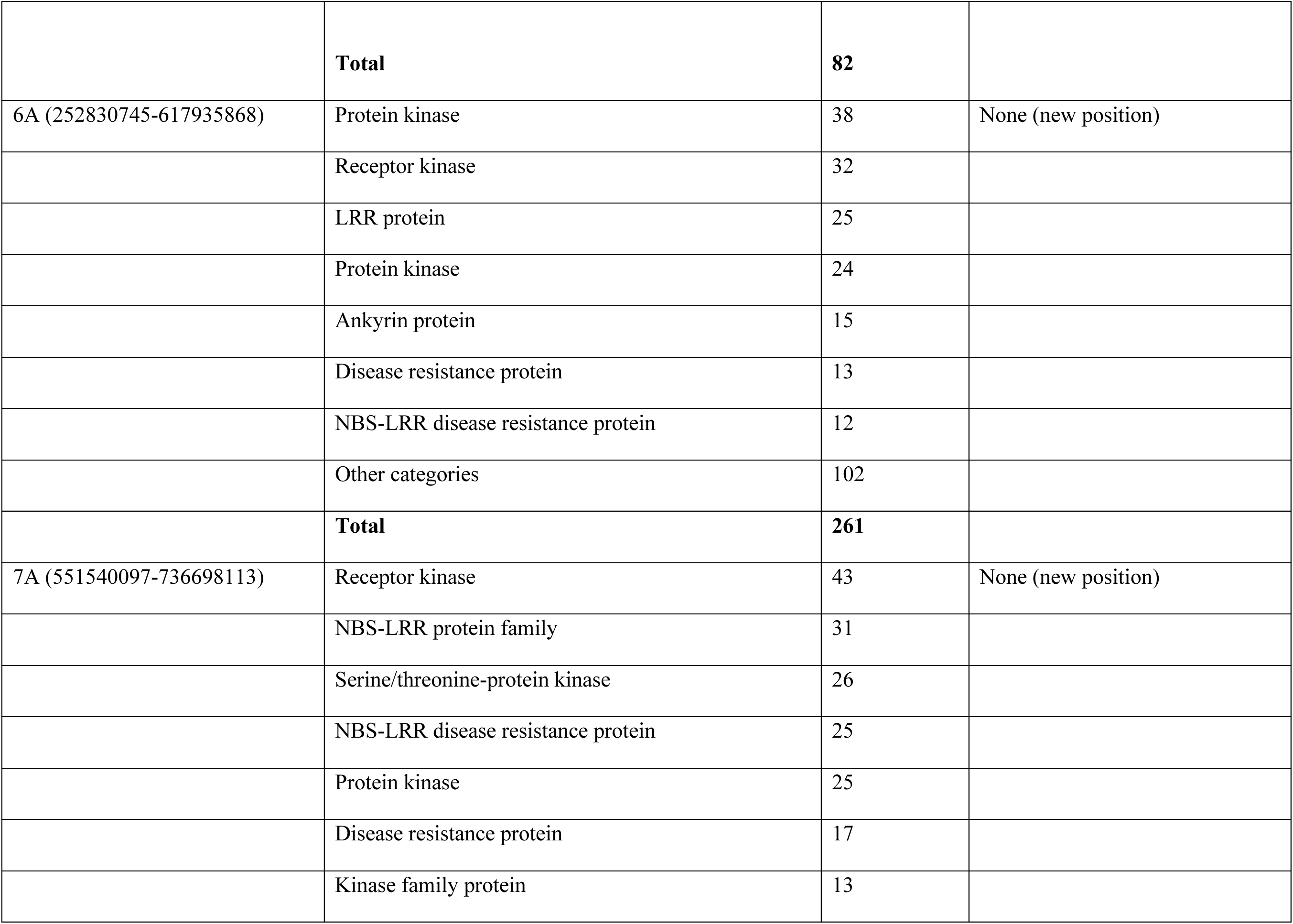
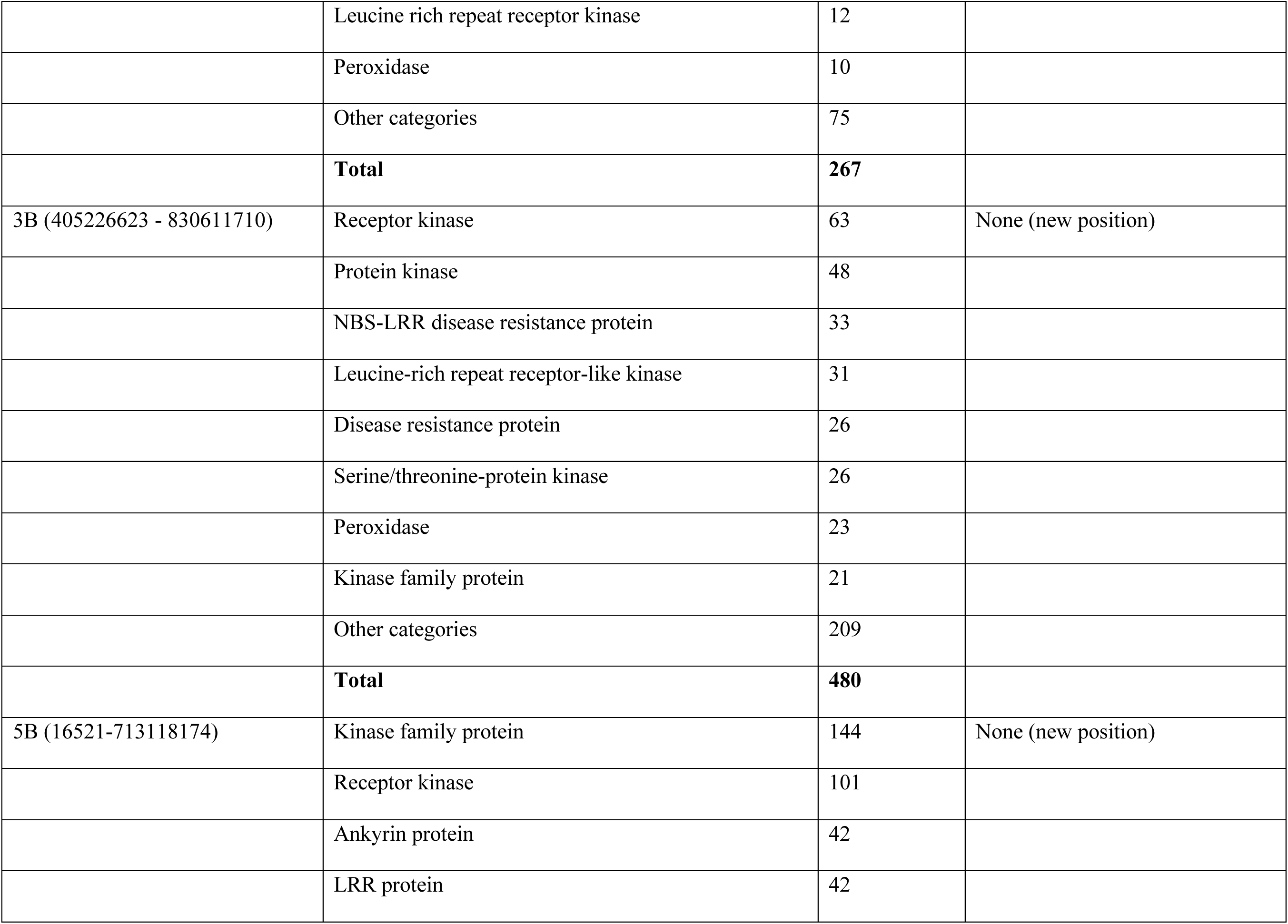
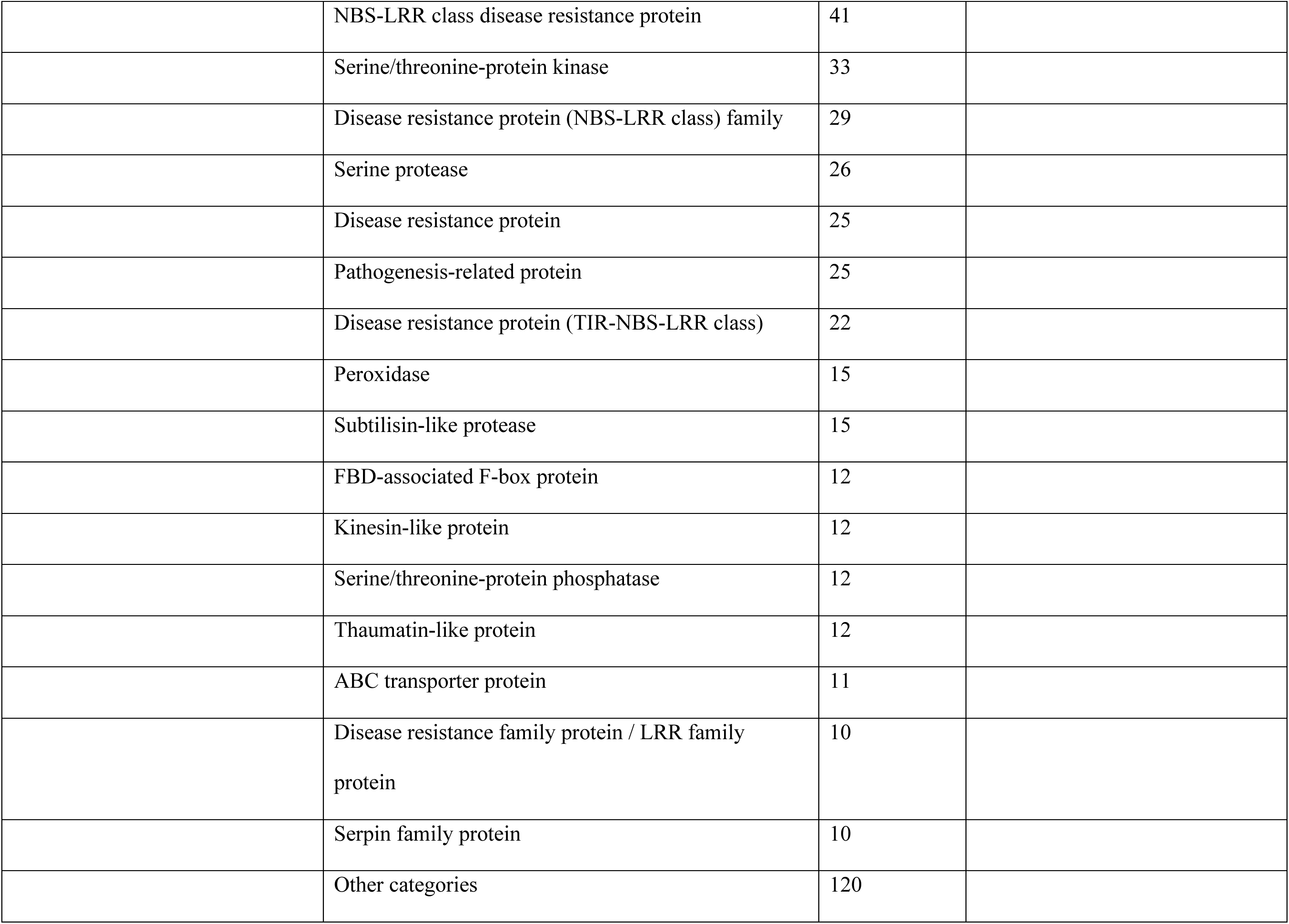
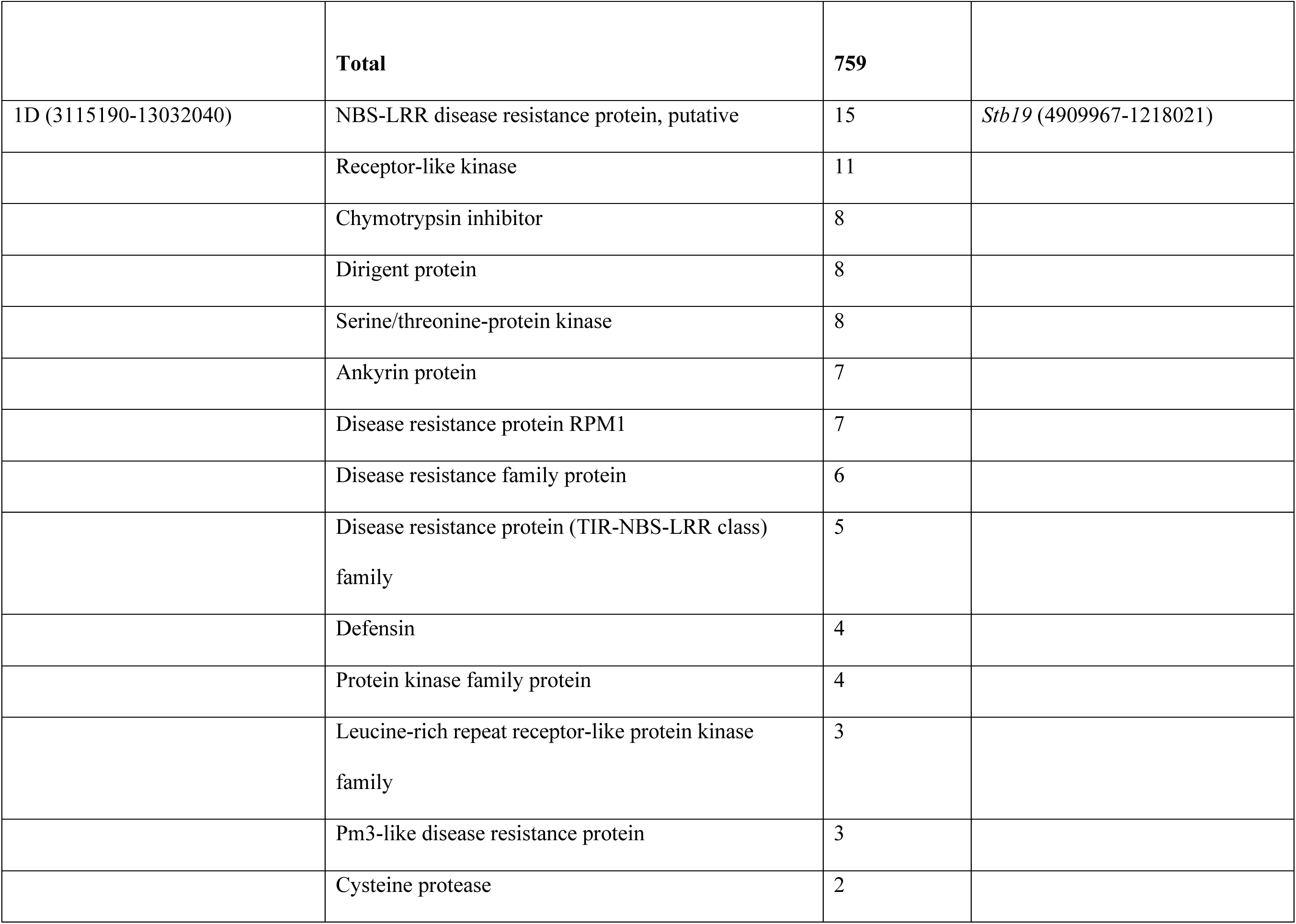
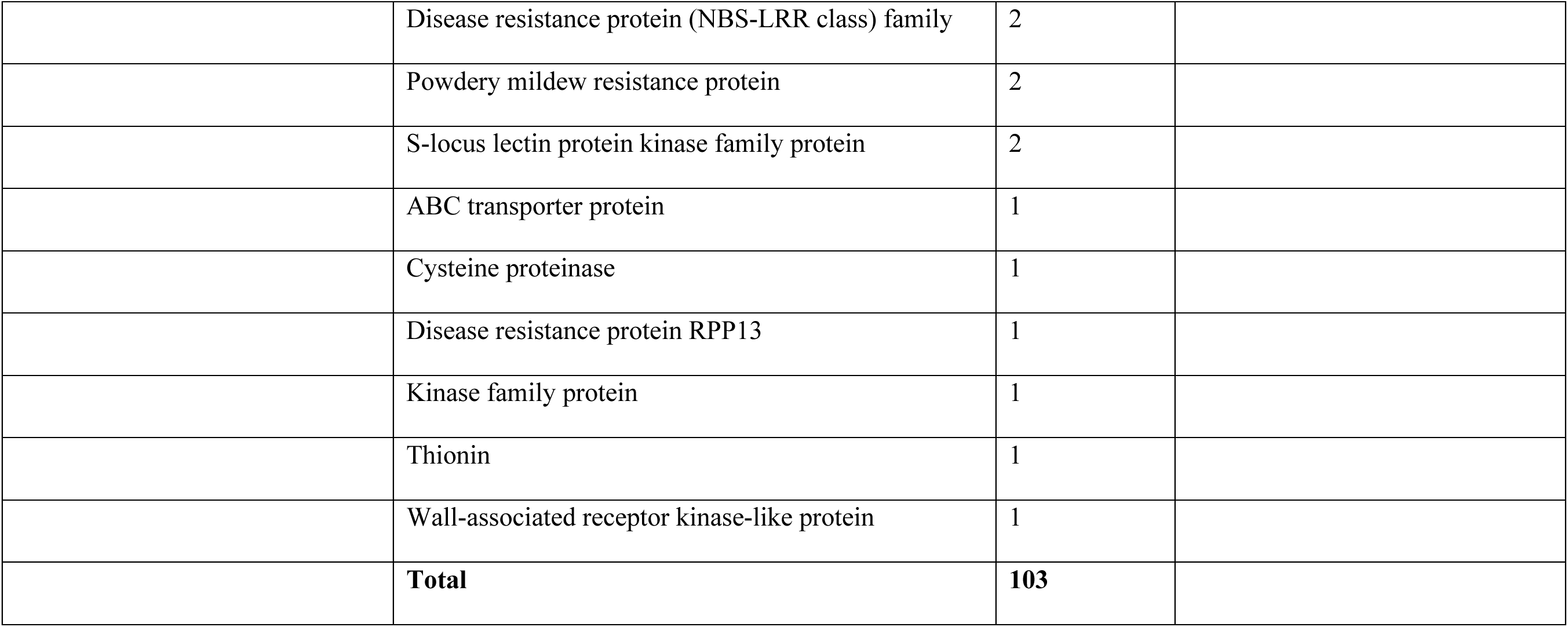
Predominant candidate resistance gene categories in genomic regions associated with resistance to *Zymoseptoria tritici* in the genome-wide association analysis.

Candidate resistance gene categories were variable from one genomic region to another, likely indicating variable pathways to express resistance to *Z. tritici*. The highest number of putative resistance genes was identified on chromosome 5B, with a total of 759 candidate genes. In contrast, only 82 candidate resistance genes were identified on chromosome 6A (20862967-75370938) (Table 7, Supplementary Tables 4 and 5).

## Discussion

Even though the soft red winter wheat panel population was composed mostly of advanced breeding lines, the GWA analysis identified significant variation for Septoria tritici blotch resistance, including putatively four previously known plus seven possibly novel loci. These genes, if confirmed, are much more likely to be effective against populations of the pathogen in the Midwestern part of the United States where most of the selection to develop the lines in this wheat panel occurred over multiple years and locations. All reported MTAs were confirmed by the FarmCPU and the CMLM (GAPIT) statistical models, with higher significance levels of the associations for the FarmCPU model compared to the CMLM model in the GAPIT program. Similar observations have been reported by Wei et al. (2017b) and Kaler et al. (2020), indicating that the FarmCPU method detected more loci and with higher levels of significance for their associated SNPs.

Different results from analyzing single-time-point scores versus AUDPC confirms that GWA results are highly contingent on the type of the inputted phenotypic data; careful consideration should be given to phenotypic data to ensure accurate GWA results (Ibrahim et al. 2020; Zhu et al. 2008). Calculation of AUDPC has been a common measure of quantitative disease resistance in plants but it entails repeated disease assessments (Jeger and Viljanen-Rollinson 2001). It has been well demonstrated that the power of detecting QTLs and MTAs is highly increased with repeated measurements (Arbelbide et al. 2006; Yu et al. 2005; Zhu et al. 2008). Therefore, AUDPC measures are potentially more accurate phenotypes for MTA discovery. Nonetheless, implementing different phenotypic variations may lead to different GWA results as confirmed by our data.

Population structure is one of the key factors that may affect resolution of a GWAS so must be considered carefully to identify true associations (Alqudah et al. 2020; Mohammadi et al. 2020; Pritchard et al. 2000). Population structure can be addressed using a statistical approach that aims to calculate relatedness correlations among individuals within the population due to admixture and/or historical structure (Alqudah et al. 2020). An unrecognized population structure will negatively impact the GWAS resolution, resulting in false-positive associations (Hayes 2013; Pritchard et al. 2000). To improve the resolution power of our GWA study, the Purdue population structure was investigated through a principal component analysis that resulted in the identification of four subpopulations. This observation agreed with that of Gaire et al. (2020) that there were four subpopulations in the soft red winter wheat panel population, consisting of two North American subpopulations and Australian and Chinese subpopulations. Structure of this population was investigated using the STRUCTURE software by Gaire et al. (2020). Both STRUCTURE (Pritchard et al. 2000) and principal component analysis (PCA) (Price et al. 2006) are common approaches that use genetic markers to determine population structure (Kaler et al. 2020). As observed in our analysis, results from STRUCTURE and PCA are usually similar (Kaler et al. 2020). However, due to lower required computational resources, PCA is generally used more commonly to generate covariates (Kaler et al. 2020). The number of sub-populations was used subsequently as a covariate in our GWAS statistical model to control for false positives that can arise from population structure and family relatedness (Kaler et al. 2020).

Two approaches were used to detect any possible false associations (Alqudah et al. 2020; Wei et al. 2017b). Application of mixed-model methods to correct for population structure using a PCA (Alqudah et al. 2016) or kinship matrix (Nagel et al. 2019) is used commonly (Alqudah et al. 2020; Crowell et al. 2016; Wallace et al. 2014). While both the FarmCPU and CMLM models control for population structure and relatedness to reduce the detection of false-negative associations (Kaler et al. 2020), these two statistical models handle the testing of markers, population structure and kinship differently (Wei et al. 2017b). The CMLM model implemented in the GAPIT package is based on the mixed linear model (MLM) and deals with large and computationally challenging datasets by clustering individuals into groups in the kinship matrix (Lipka et al. 2012). Hence, this model fits genetic values of groups as random effects in the model, thus reducing false negatives and enhancing the identification of true associations (Kaler et al. 2020).

The FarmCPU model is based on a Multiple Loci linear Mixed Model (MLMM) that incorporates multiple markers simultaneously as covariates in a stepwise Mixed Linear Model (MLM) to partially remove the confounding among markers that occurs with kinship (Kaler et al. 2020; Liu et al. 2016b). Compared to CMLM, in which the kinship matrix remains constant for all the markers, FarmCPU adjusts its kinship matrix based on the tested markers and covariates in the fixed-effects model (Wei et al. 2017b). Another advantage of the FarmCPU model compared to CMLM is the use of selected associated markers as covariates and the simultaneous testing of multiple markers, which significantly improves the control of both false-positive and false-negative associations (Liu et al. 2016b). Therefore, despite the popular use of the CMLM model for GWAS, FarmCPU provides for better control of the confounding among population structure, kinship, and the simultaneous testing of markers, leading to an increased statistical power with fewer false positives (Liu et al. 2016b; Wei et al. 2017b). Nonetheless, it is always advisable and beneficial to implement different models to test for associations to increase the confidence in the overlapping loci, and to carefully choose the best approach to conduct GWAS based on the species and traits being analyzed (Wei et al. 2017b).

Another potential stumbling block for identifying resistance genes is selection of the most appropriate isolate(s) for phenotypic testing. As cultivar-specific interactions are observed commonly between *Z. tritici* and wheat (Ahmed et al. 1995; Brown et al. 2001; Cowger et al. 2000; Kelm et al. 2012; Kema et al. 1996a; Kema et al. 1996b; Kema et al. 2018; Kema and van Silfhout 1997; Kema et al. 2000), a careful selection of the isolates for testing is essential for successful identification and postulation of genes (Adhikari et al 2004; Ghaffary 2011; Ghaffary et al. 2018). To address this issue, a seedling pre-screening assay was conducted on a suite of differential bread wheat cultivars harboring known *Stb* major resistance genes to identify the best testers among *Z. tritici* isolates sampled from diverse U.S. regions. This assay revealed a variable pathogenicity pattern, particularly for pycnidia development, which confirmed the specific cultivar × *Z. tritici* interaction in the wheat–*Z. tritici* pathosystem, thus indicating that specific gene actions govern STB resistance. This pre-screening enabled us to select two *Z. tritici* isolates that were used subsequently to evaluate seedlings of the soft red winter wheat panel population.

These preliminary tests identified pycnidia development as the most discriminating trait among all tested isolates compared to necrosis development. Hence, pycnidia development appears to be the most accurate phenotypic criterion to use for isolate selection with our materials, and also has been shown to give the best discrimination among resistance genes in other *Z. tritici*-wheat interaction analyses (Dalvand et al. 2016; Ghaffary 2011; Ghaffary et al. 2018; Kema et al. 1996a; Kema et al. 1996b; Kema and van Silfhout 1997). Even so, the subsequent seedling assays revealed specific interactions for both necrosis and pycnidia development. Therefore, the tested population appears to harbor divergent resistance loci that are specific for each isolate, with necrosis and pycnidia development likely under different genetic control. This result agrees with previous demonstrations that necrosis and pycnidia resistance are controlled by divergent genetic factors (Stewart et al. 2016; Stewart and McDonald 2014).

Comparisons with genomic locations of known major *Stb* resistance genes revealed that, irrespective to the phenotypic datasets used, the GWA analysis unveiled multiple novel locations associated with likely *Z. tritici* resistance QTL and others that corresponded to previously mapped genes. While AUDPC measures generated with the T2 isolate of *Z. tritici* indicated that the soft red winter wheat panel lines probably contain the major gene *Stb19* (Yang et al. 2018), novel locations for STB resistance were detected when using single time-point scores generated by the same isolate, on chromosomes 3A (*S3A_699078407*), 3B (*S3B_736712248*), 4B (*S4B_660493440*), 5A (*S5A_666525700*), 6A (*S6A_14386770*), 6D (*S6D_447068059*), 7A (*S7A_38234502*; *S7A_38238713*), and 7B (*S7B_11882709*, *S7B_605313385*, *S7B_711755909*), as well as the MTAs *S7A_84027035*, *S7A_85912163*, and *S7A_91722899* on chromosome 7A that overlapped with the location of the known major resistance gene *Stb3* (Goodwin and Thompson 2011; Goodwin et al 2015).

Novel genomic locations also were revealed by the T46 AUDPC scores on chromosomes 1A (two locations: *S1A_500074551;* and *S1A_577377232*), 3B (*S3B_726510291*), 4A (*S4A_605028654*), 6A (*S6A_532657432*), and 7A (*S7A_719723711*) for pycnidia and 6A (*S6A_611714334*) for necrosis. These MTAs were confirmed using the single-time-point scores that additionally detected the new genomic positions on chromosomes 2D (S2D_29753796), 4B (S4B_548241937) and 4D (S4D_439658613).

Previous analyses also identified markers on chromosomes 1A, 3A, 5A, 6A, 7A, 3B, 4B, 5B, 7B, 2D, 4D and 6D that were associated with STB resistance (Gurung et al. 2014; Muqaddasi et al. 2019a; Odilbekov et al. 2019; Vagndorf et al. 2017; Yates et al. 2019). However, we could not ascertain whether these positions overlap with our reported locations due to the lack of the physical position information and /or the sequence data of the previously reported MTAs. Hence, the only accurate and possible comparison was towards the major known *Stb* resistance genes.

Locating the associated SNPs on the bread wheat genome sequence IWGSC RefSec v1.0 allowed candidate resistance genes to be identified within the LD blocks of the MTAs on each chromosome. Many of the candidate genes that were identified encode classes of proteins known to be involved in plant disease resistance pathways (Supplementary Tables 4 and 5). By far the largest class encodes proteins with leucine-rich repeat domains and nucleotide-binding domains. NB-LRR proteins are frequently identified as conferring gene-for-gene resistance (Eitas and Dangl, 2010). Mechanistically, they interact either directly or indirectly with effector proteins produced by avirulent pathogens, triggering a signaling cascade that ultimately causes cell death. The high frequency of NB-LRR genes being associated with STB resistance loci is consistent with the fact that this type of gene is very often found in clusters that undergo frequent recombination and sometimes generate new R-gene specificities (McHale et al., 2006). In addition, many receptor kinases were found (Table 7), which is consistent with several recently cloned *Stb* genes (Hafeez et al 2025; Saintenac et al 2018; Saintenac et al 2021), including wall-associated receptor kinases like *Stb6* (Saintenac et al 2018) and lectin receptor kinases (Supplementary Table 4) that may be similar to *Stb15* (Hafeez et al 2025). Our analyses were not able to discern which of these genes might have essential functions in STB resistance but did identify numerous candidates for future research. This can include functional analyses, such as loss of function studies with mutants, gene editing, or expression knockdown by RNAi or Virus-Induced Gene Silencing (Scofield et al 2006).

In conclusion, our work shed light into the genetic make-up of septoria tritici blotch resistance factors in elite breeding lines of U.S. soft winter wheat. Known and novel genomic regions were identified. Additional work is required to validate the novel QTL and narrow down their genomic locations. However, once carefully validated these potentially novel QTL could be used by wheat breeding programs to increase the level of resistance to STB in the future.

## Supplementary Information

The online version contains supplementary material available at URL.

## Acknowledgements

The authors thank Ian Thompson for locating and helping acquire space to perform the inoculations.

## Author contributions

LA mostly designed the experiments, performed the analyses and wrote the first draft of the manuscript. RG helped with data analysis. GB-G supervised generation of the molecular genotyping data. SS provided funding and edited the manuscript. MM hired LA to do the work, helped with analyses and with improving the manuscript. SBG helped design the experiments, provided funding, wrote parts and edited the entire manuscript.

## Data availability

The datasets generated and/or analyzed during this research are available from the corresponding author on reasonable request.

## Supplementary information

**Supplementary Table 1**. Phenotypic response at 21 days post inoculation, in percentage necrosis and pycnidia on the inoculated primary leaves, of differential lines to 13 *Zymoseptoria tritici* isolates. Colored cells indicate significant differences (LSDs; P=0.05) with resistant in green (not significantly different from 0 %), intermediate in yellow (significantly different from 0 % and 100%) and susceptible in red (not significantly different from 100%).

**Supplementary Table 2**. Analysis of variance of the necrosis and pycnidia scores on seedlings of the differential wheat set tested with the T2 and T46 isolates of *Zymospetoria tritici*.

**Supplementary Table 3**. Marker-trait associations with STB resistance mapped on the soft red winter wheat panel population with the T2 and the T46 isolates of *Zymoseptoria tritici* at the single time point score (16, 18 and 22 days-post inoculation) using models from FarmCPU and GAPIT (CMLM).

**Supplementary Table 4.** List of candidate resistance genes identified using the wheat reference genome IWGSC RefSec v1.0.

**Supplementary Table 5.** Resistance candidate gene IDs and coordinates.

**Supplementary Figure 1**. Box plots of the Necrosis (panel **A**) and Pycnidia (panel **B**) AUDPC scores showing significant differences between the *Zymoseptoria tritici* isolates T2 and T46 when tested on the soft red winter wheat panel population

**Supplementary Figure 2**. Manhattan plots and Q-Q plots of the GWAS analysis generated by the GAPIT MCLM model

**Supplementary Figure 3**. Categories for candidate genes identified in the indicated genomic regions associated with resistance against *Zymoseptoria tritici* from the Genome-Wide Association analysis

