## Supplementary Fig. S1 for "Genome-Wide Association study in a US soft winter wheat population reveals novel and known sources of resistance to the Septoria tritici blotch pathogen *Zymoseptoria tritici*"

### Slide 1
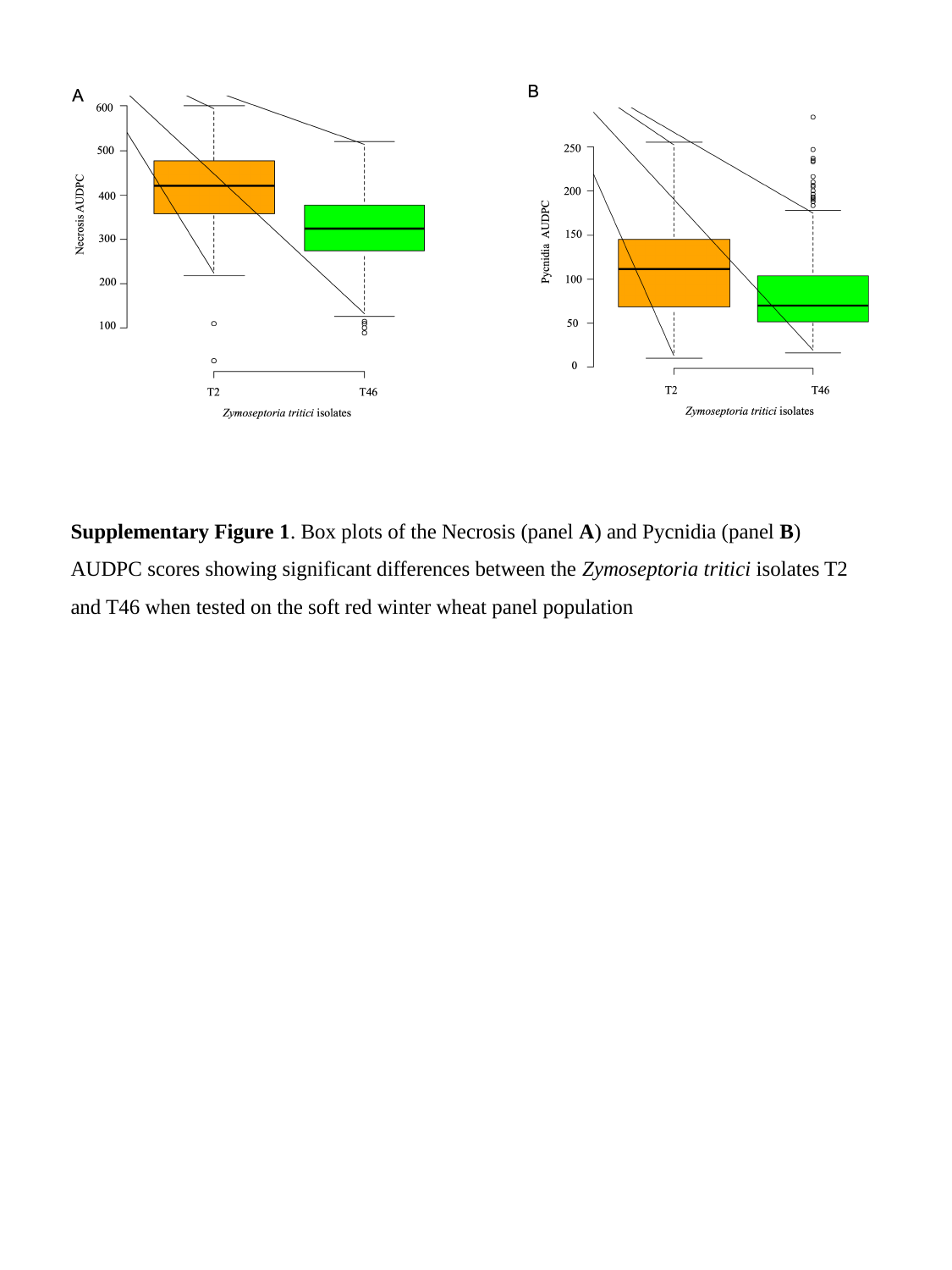

Supplementary Figure 1. Box plots of the Necrosis (panel A) and Pycnidia (panel B) AUDPC scores showing significant differences between the Zymoseptoria tritici isolates T2 and T46 when tested on the soft red winter wheat panel population
