## Supplementary Fig. S2 for "Genome-Wide Association study in a US soft winter wheat population reveals novel and known sources of resistance to the Septoria tritici blotch pathogen *Zymoseptoria tritici*"

### Slide 1
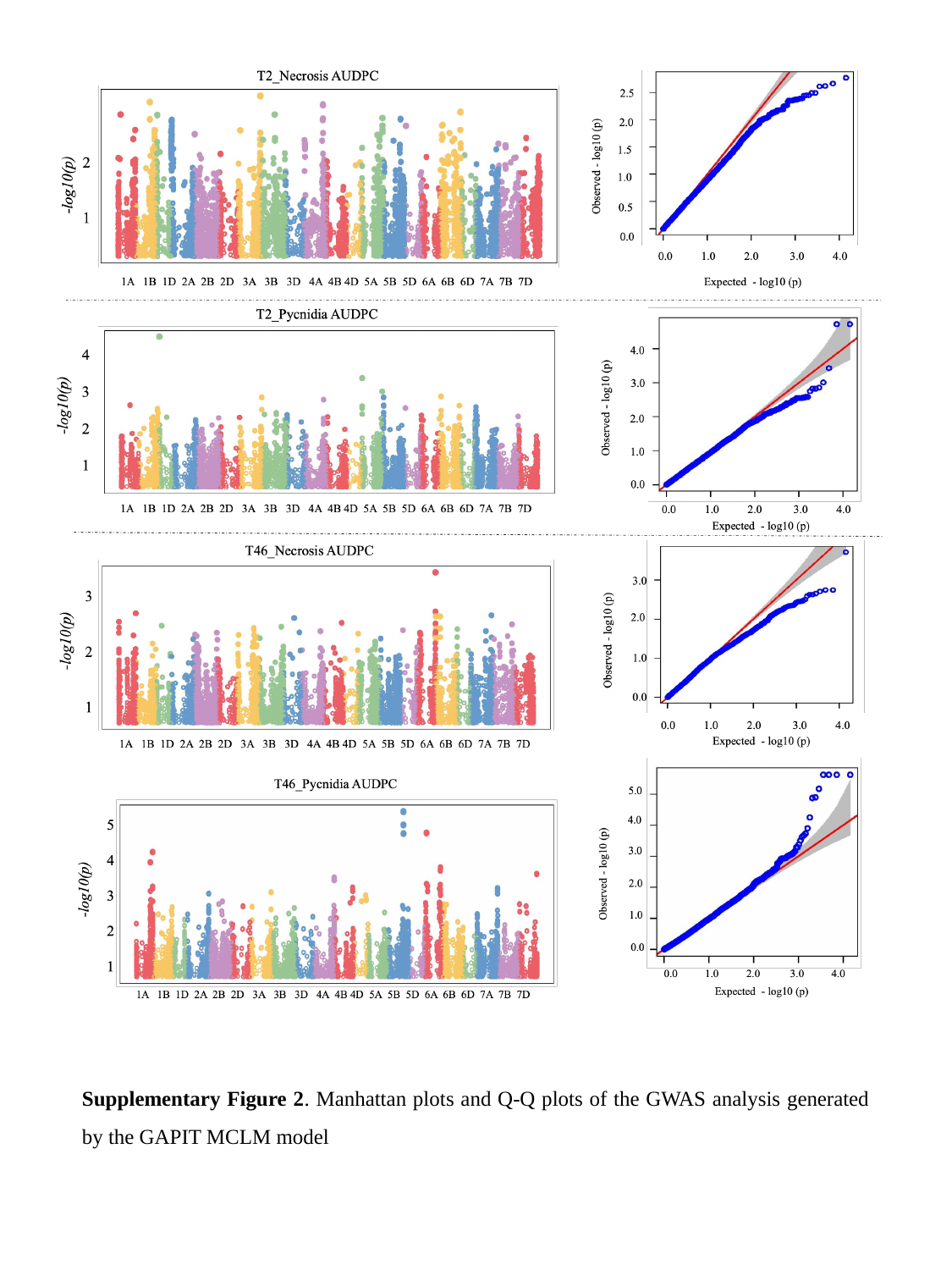

Supplementary Figure 2. Manhattan plots and Q-Q plots of the GWAS analysis generated by the GAPIT MCLM model
