## Supplementary Fig. S3 for "Genome-Wide Association study in a US soft winter wheat population reveals novel and known sources of resistance to the Septoria tritici blotch pathogen *Zymoseptoria tritici*"

### Slide 1
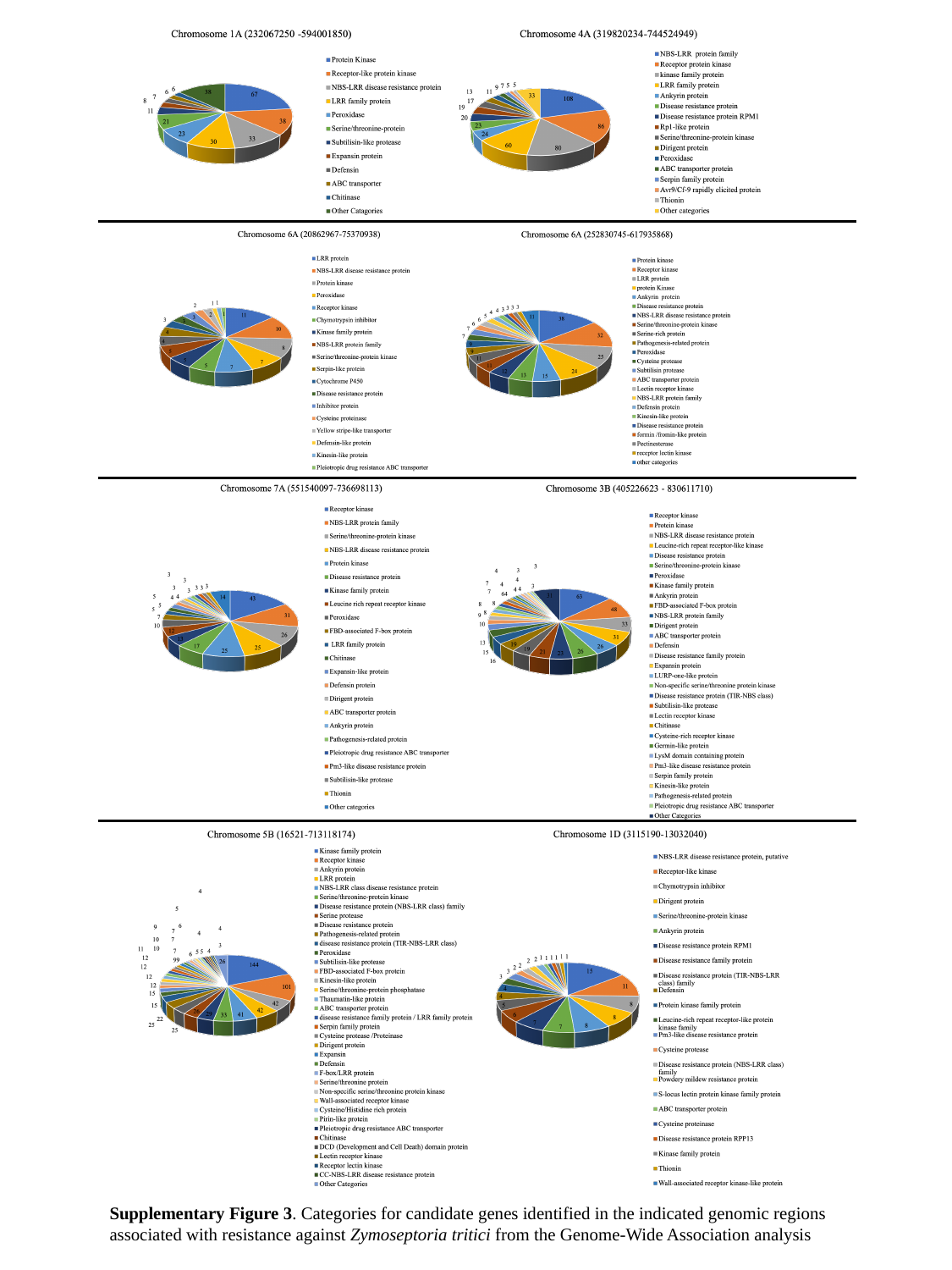

Supplementary Figure 3. Categories for candidate genes identified in the indicated genomic regions associated with resistance against Zymoseptoria tritici from the Genome-Wide Association analysis
